# The RNA-binding protein Imp1 promotes a Spdef transcriptional program and mucus fucosylation during necrotizing enterocolitis

**DOI:** 10.64898/2026.01.30.702645

**Authors:** Kevin A. Swift, Alexandria J. Shumway, Molly Aloia, Madeline Hedges, Rachel Pung, Cariam Rodriguez Santiago, Michael Shanahan, Alexander Drake, Melanie H. Hakar, Leigh Selesner, Madeline Kuhn, Claire Yung, Praveen Sethupathy, Sarah F. Andres

## Abstract

**Background:** In the United States over 10% of all neonates are born premature (less than 37 weeks gestational age), and many face complications related to prematurity, including necrotizing enterocolitis (NEC). NEC is the most deadly gastrointestinal disease and the leading cause of death in preterm neonates, with up to 50% mortality. Since there is no cure for NEC, prevention is the best strategy. Enhancing our understanding of intestinal epithelial cell (IEC) responses to NEC damage will provide novel therapeutic targets to prevent NEC.

Published evidence suggests that the RNA-binding protein insulin-like growth factor 2 mRNA binding protein 1 (IMP1) plays roles in intestinal development, barrier function, and intestinal repair. Notably, however, roles for IMP1 in NEC are not defined. Goblet cells produce protective mucus in the intestine, and their mature function is dependent on the transcription factor Spdef. Emerging evidence suggests that goblet cell mucus complexity impacts barrier function and inflammation susceptibility. This study aimed to define the role of IMP1 in NEC pathogenesis using neonatal human enteroids and a model of NEC-like intestinal injury in mice with IEC-specific Imp1 overexpression and loss.

**Hypothesis:** IMP1 expression is protective in NEC.

**Methods:** This study used mice with intestinal epithelial Imp1 overexpression or loss and corresponding wild-type controls. At post-natal day 3, mice of both sexes were randomly assigned to control or NEC groups. NEC was induced with the well-established experimental NEC-like intestinal injury model that includes stress, formula feeding, and hypoxia. Imp1 effects on experimental NEC were assessed using RNA sequencing, western blotting, and immunostaining.

**Results:** Inflammatory bacteria induced *IMP1* expression in neonatal human enteroids. Mice with Imp1 overexpression incur worse intestinal damage during NEC. Pathway analysis of RNA sequencing data revealed a significant enrichment of the Spdef transcriptional network in *Imp1^IEC-OE^* during NEC. This included significant upregulation of Spdef target genes such as *Agr2* (in NEC WT: 617.8 ± 33.56 vs *Imp1^IEC-OE^*: 812.9 ± 111.3, *p*=0.02) and *Fut2* (in NEC WT: 219.4 ± 34 vs *Imp1^IEC-OE^*: 396.6 ± 62.9, *p*=0.05). *In silico* analysis predicted Imp1 binding to *Spdef* and mucus glycosylation mediator mRNAs. Although genotype did not affect goblet cell number, *Imp1^IEC-OE^* mice with NEC exhibited significant increases (*p*0.05) in Spdef protein, genes responsible for goblet cell function (*Spink4, Klk1*, *Tspg1*) and mucus glycosylation (*Gcnt3*, *B3gnt7, Qsox1*). Ultimately, Imp1 overexpression led to increased mucus fucosylation during NEC.

**Conclusion:** Our data indicate that during NEC, upregulation of Imp1 promotes goblet cell function via Spdef, including enhanced goblet cell maturation and mucus fucosylation.

**NEW AND NOTEWORTHY:** Expression of the RNA-binding protein Imp1 is enhanced in neonatal human enteroids in response to inflammatory bacteria. Imp1 regulates goblet cell function in response to early intestinal inflammation in a mouse model of necrotizing enterocolitis. Specifically, Imp1 promotes a Spdef transcriptional program by upregulating Spdef, resulting in elevated gene expression of Spdef target genes. Additionally, Imp1 increases the gene expression of α(1,2)fucosyltransferase, *Fut2*, and subsequently, production of fucosylated mucus marked by UEA1.

**Graphical Abstract:** 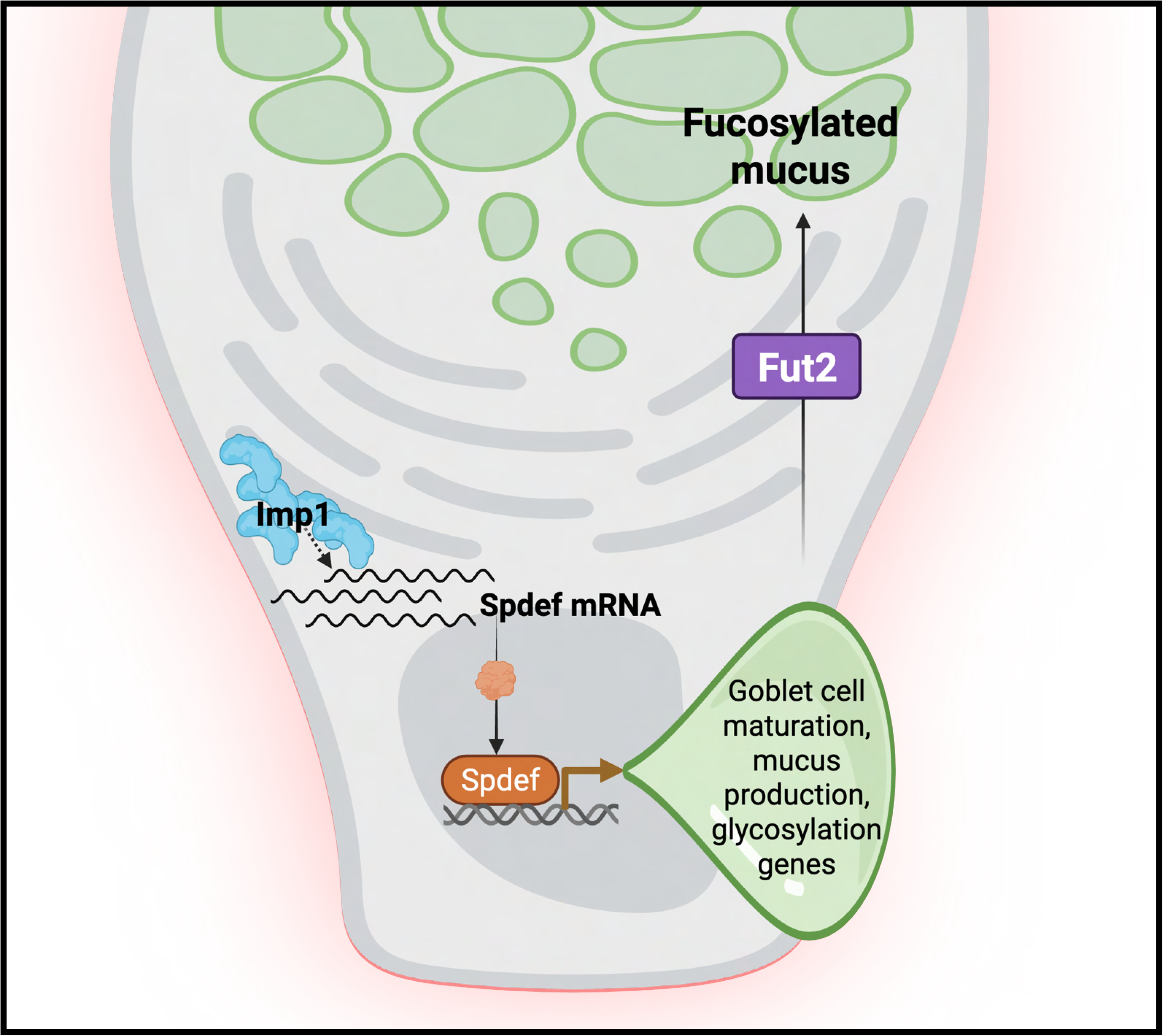

## INTRODUCTION

Necrotizing enterocolitis (NEC) is a devastating gastrointestinal disease that primarily affects very low birth weight (VLBW; <1500 grams at birth) preterm (<37 weeks gestational age) infants. In preterm infants, the disease is thought to arise from an interplay of many factors, including intestinal immaturity, dysregulated immune responses, and microbial dysbiosis. Despite advancements in neonatal care, NEC remains a significant cause of morbidity and mortality among preterm infants, highlighting the urgent need for innovative therapeutic and preventative strategies. Advancing our understanding of NEC’s molecular and cellular mechanisms is critical to improving outcomes for these vulnerable patients.

Although numerous risk factors for NEC have been identified, its precise molecular drivers remain incompletely understood. The insulin-like growth factor II mRNA-binding protein (IMP1; also known as IGF2B1, CRD-BP, ZBP1, VICKZ1) is a regulator of embryonic intestinal development (1–6) and is expressed highly in the murine intestinal epithelium until postnatal day 12 (1). Mice deficient in Imp1 exhibit severe developmental abnormalities, including underdeveloped intestines characterized by reduced villus size in the small intestine and short, irregular crypts in the colon (1). Beyond its developmental role, IMP1 expression can be up-regulated in response to damage (7) and during cancer (8–11). IMP1 is implicated in intestinal epithelial homeostasis in adult mice where it can be beneficial (12) or detrimental (7) in adult mouse models of colitis. This underscores the highly context-dependent and sometimes paradoxical role of Imp1 in disease.

Imp1 contains two RNA recognition motifs and four KH domains which govern its binding affinity and specificity for numerous targets (13). Evidence from genome-wide cross-linking and immunoprecipitation studies shows that Imp1 binds numerous transcripts through targeting the coding sequence, as well as the 3’ and 5’ untranslated regions (UTR) (14, 15). IMP1 binding plays a central role in post-transcriptional gene regulation by modulating mRNA localization, stabilization, and translation (1, 7, 16–21).

Small intestinal epithelial integrity and susceptibility to inflammatory insults, such as NEC, are strongly influenced by a layer of secreted mucus (22). Disruption of the mucus barrier in inflammatory conditions such as colitis can lead to dysfunctions associated with NEC, including intestinal barrier leakiness, infection, and inflammation (23–25). The mucus barrier is secreted by intestinal epithelial goblet cells, serving both as a physical defense against pathogens and a nutrient source for commensal microorganisms (22). Mucins, the main constituents of mucus, undergo extensive glycosylation, which enhances their barrier properties and selects for specific intestinal microbes. For example, fucosylation or the addition of fucose moieties to intestinal mucus enhances barrier function and promotes the expansion of beneficial commensal bacteria (26, 27).

The role of Imp1 in early intestinal responses to inflammatory damage, such as NEC, is unknown. Therefore, we used neonatal human enteroids and genetic mouse models with Imp1 loss or overexpression to examine the role of intestinal epithelial Imp1 during early life inflammation and tested the hypothesis that Imp1 expression would be protective against inflammatory insult.

## METHODS

### Neonatal human enteroid generation and culture

Collection of tissue samples used for enteroid generation was approved under OHSU IRB # 21952. Intestinal enteroid lines were established from neonatal intestinal tissue as described previously (28)Frazer, 2023 #100}. Demographic details of participant samples are shown in Table 1. Enteroids were maintained and propagated in Matrigel (Growth factor reduced organioid culture 356255 and 354234) patties in the ‘basal out’ orientation using IntestiCult Growth Medium (STEMCELL Technologies, Vancouver, Canada).

**Table 1.** Intestinal tissue sample demographics.

| ID | Sex | Gestational Age | Corrected Gestational Age at Tissue Collection | Sample Location | Diagnosis |
| --- | --- | --- | --- | --- | --- |
| 4 |  | 39 |  |  | Ileal Duplication Cyst |
| 19 | M | 23 6/7 | 44 6/7 | Jejunum | Prior NEC, Jejunostomy Reversal |
| 25 | F | 34 1/7 | 34 5/7 | Proximal Jejunum | Jejunal Atresia |
| 26 | M | 36 4/7 | 51 4/7 | Ileum | Gastroschisis, Ileostomy Reversal |

### Apical-out enteroid- NEC-in-a-dish model

To gain access to the apical/luminal surface of human enteroids, basal-out enteroids were everted to an apical-out orientation via polarity reversal. Apical-out eversion was performed as previously described and validated (28). After 48-hours in suspension culture, enteroids were treated for 8 hours with a 1:100 dilution of a UV-killed enteric bacteria (1.23 x 10^8^ CFU/mL) isolated from an infant with NEC totalis or PBS as a control. The mixed bacterial culture contains predominantly Proteobacteria and Enterobacteriaceae and was originally described by Good et al (29). After 8 hours of stimulation, enteroids were collected for RNA isolation and qRT-PCR, as described below.

### Imp1 mouse models and breeding

All animal studies were approved by the Institutional Animal Care and Use Committee at the Oregon Health and Science University (OHSU) under protocol #3036. Mice were bred in a super-barrier facility at OHSU. Mice lacking Imp1 in the intestinal epithelium, *VillinCre;Imp1^fl/fl^*and called “Imp1^ΔIEC^” mice, were obtained by mating *VillinCre* mice (C57BL/6J, Jackson Laboratories, Bar Harbor, ME USA 04609) with *Imp1^fl/fl^* mice generously provided by Dr. Vladimir Spieglman and described previously(9). Mice with Imp1 overexpression in the intestinal epithelium, *VillinCre;lsl-FLAG-Imp1* and called “Imp1^IEC-OE^” mice were obtained by mating *VillinCre* mice with mice containing Flag-Imp1 cDNA targeted to the Rosa locus downstream of a polyA sequence flanked by loxP sites (lox-STOP-lox, lsl), generously provided by Dr. Kate Hamilton, to selectively overexpress Imp1 in the intestinal epithelium, as originally described(9). All genotyping was performed by Transnetyx using validated primer probe sets.

### Intestinal epithelial cell (IEC) Isolation

Intestinal segments were dissected from p6 mice and kept in ice-cold DMEM high glucose medium (Gibco, ThermoFisher, Waltham, MA, USA) until processing. Segment length for each isolation is as follows. Mass-Spectrometry Proteomics: Entire Small bowel. Western, qPCR Genotyping: 6 cm ileum just proximal to the cecum. These segments were splayed open to expose the epithelial surface.

Intestinal segments were vortexed 4 times (15 seconds vortex, 15 seconds on ice) in Intestinal Wash Solution (IWS): calcium- magnesium free Hanks’ Balanced Salt Solution (CMF-HBSS) (Gibco, ThermoFisher) with 1 mM N-acetyl cysteine (NAC) (Millipore Sigma Burlington, MA.). Samples were pelleted at 300 x g and transferred into IWS with 10 mM ethylenediaminetetraacetic acid (EDTA)(Invtrogen, ThermoFisher). . Samples were then incubated on a rotator at 4°C for 45 minutes. Following incubation, samples were vortexed 4 times for (30 seconds vortex, 30 seconds on ice), to lift IECs from the basement membrane.

Muscle, basal lamina, and mesenchyme pieces were then removed using forceps. IECs were pelleted at 300 x g, washed with 1x PBS 1 time and either immediately prepared as described below, or snap-frozen for future use.

### Mouse model of NEC-like intestinal injury

The NEC model was established following previously described methods, with minor modifications (30) (Figure 1A). Three-day-old neonatal mice (P3) were randomly assigned to either the control or experimental NEC group. Littermate, control mice remained with the dam and continued to nurse (“dam-fed”). To induce NEC, pups were separated from the dam, housed in an infant incubator at 37 °C (OHMEDA Care Plus Incubator; Ohio Care, Norcross, GA), orally gavage-fed a formula mixture, and exposed to hypoxic conditions twice daily. The formula feed was a combination of Similac Advance Infant Formula (Abbott Nutrition, Columbus, OH) and Puppy Milk Replacer (ESBILAC; PetAg) supplemented with lipopolysaccharide (LPS) and enteric bacteria isolated from an infant with NEC totalis (29, 30). LPS was added at a dose of 3 µg/g for litters with an average pup weight below 2 g and 4 µg/g for litters averaging above 2 g. NEC pups were gavage-fed using a 1.9-French single-lumen silicone peripherally inserted central catheter (PICC-Nate; Utah Medical Products, Midvale, UT) every 3 hours between 7:30am – 10pm over the course of 72 hours beginning at 10 am on p3 and concluding at 10pm on p5. A. To induce hypoxia, NEC pups were placed in a modular chamber (BioSpherix, Parish, New York) containing 5% O₂ and 95% N₂ for 10 minutes, twice daily, over the three-day NEC induction. All mice weighed each morning. On postnatal day 6, all pups were euthanized for tissue collection in accordance with IACUC standards. The intestine was isolated, placed in ice-chilled phosphate-buffered saline (PBS), linearized, and the mesentery was removed with forceps and scissors. Half-centimeter segments of terminal ileum were collected retrograde from the cecum as shown (Figure 1B).

**Figure 1.**
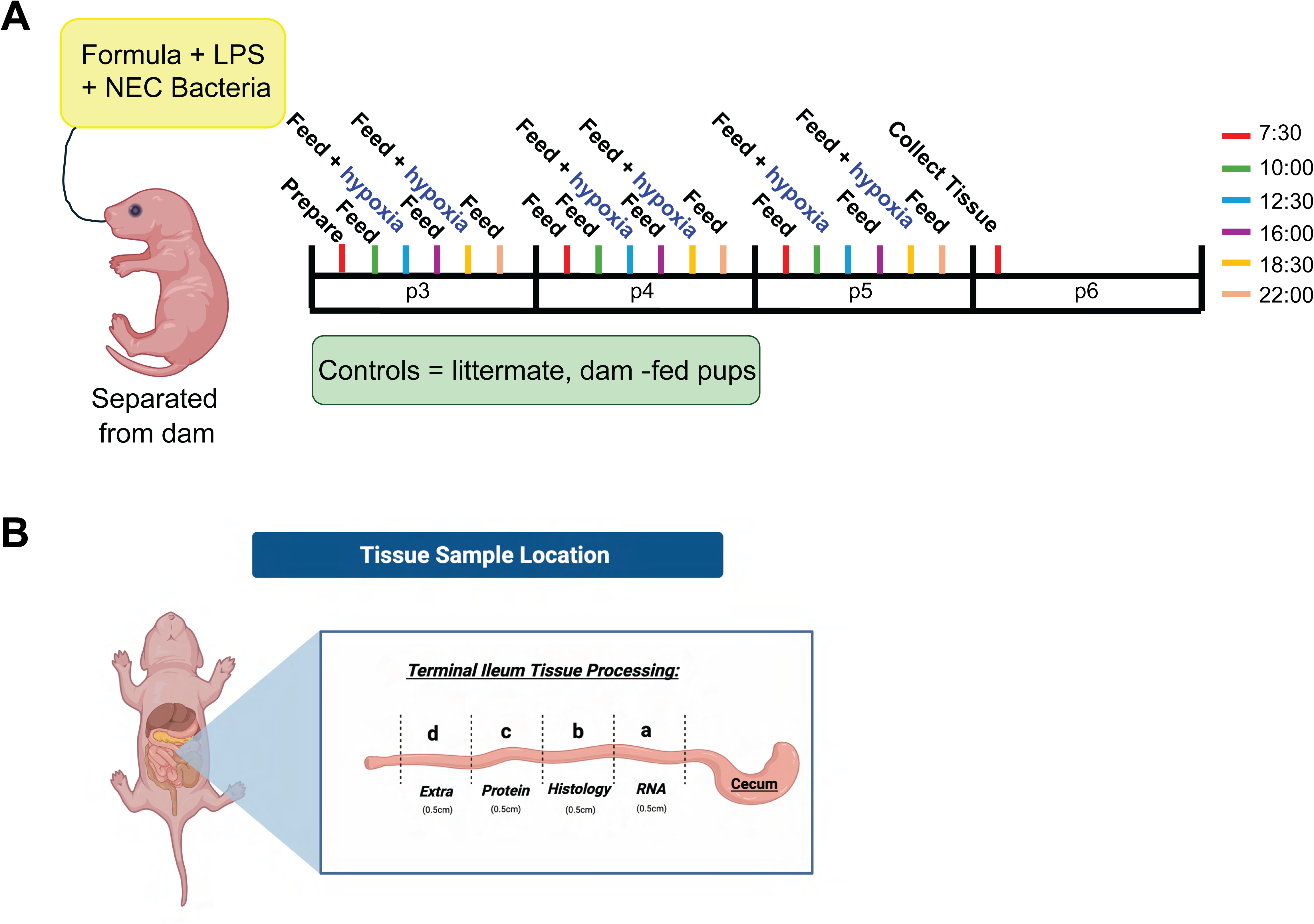
Mouse model of NEC-like intestinal injury. A) NEC-like intestinal injury was induced in post-natal day 3 (p3) mice. Mice were randomly assigned to either the control or the experimental NEC group. Littermate, control mice remained with the dam and continued to nurse (“dam-fed”). To induce NEC, pups were separated from the dam, housed in an infant incubator at 37°C, orally gavage-fed a formula mixture of human infant and canine formula laced with lipopolysaccharide (LPS) and enteric bacteria isolated from an infant with NEC totalis¹, and exposed to hypoxic conditions (5% O₂) twice daily for three days. At the conclusion of the experiment (post-natal day 6), B) Terminal ileal tissue was harvested and allocated for the following downstream analyses: (a) RNA isolation, (b) fixation for histology, (c) protein isolation, (d) additional analysis as needed.

### Whole thickness tissue preparation for Protein or RNA Extraction

Whole-thickness intestinal segments (0.5 cm) were lysed with 1.0 mm zirconia/silica beads (BioSpec Products, Bartlesville, OK, USA) in a mini-bead beater (BioSpec Products). Samples were lysed in radioimmunoprecipitation assay-sodium dodecyl sulphate (RIPA-SDS)(Cell Signaling Technologies, Danvers MA) for western blot or GeneJet RNA Purification Lysis Buffer (Thermo Fisher) for RNA purification. Samples in RIPA-SDS were frozen for future use, while samples lysed for RNA were immediately prepared.

### RNA Purification, cDNA Synthesis, and RT-qPCR

#### Enteroids

Total RNA was isolated using the RNAqueous Micro Total RNA Isolation Kit (Thermo Fisher Scientific) according to the manufacturer’s protocol and including an on-column, DNase treatment step (Qiagen, Valencia, CA). RNA concentration was determined using 260 nm absorbance with a Nanodrop Spectrophotometer (Thermo Fisher), and quality was assessed by RNA integrity gel (1% agarose), when RNA yield allowed.

### Mouse tissue

RNA was purified using GeneJET RNA Purification Kit (Thermo Fisher) according to the manufacturer’s protocol and including an on-column, DNase treatment step (Qiagen). RNA concentration was determined using 260 nm absorbance with a Nanodrop Spectrophotometer (Thermo Fisher), and quality was assessed by RNA integrity gel (1% agarose). RNA was used for RNA sequencing (described below) and cDNA was synthesized for qPCR using High-Capacity cDNA Reverse Transcription Kit (Applied Biosystems, Thermo Fisher) and a C1000 Touch™ thermocycler. RT-qPCR was performed with TaqMan Fast Advanced Master Mix reagents (Advanced Biosystems, Thermo Fisher) according to manufacturer’s protocol. qPCR was performed with TaqMan reagents and StepOnePlus (Advanced Biosystems, Thermo Fisher) PCR systems using the following probes: <u>human:</u> ribosomal protein lateral stalk subunit P0 (RPLP0, Hs99999902_m1) and insulin-like growth factor 2 mRNA-binding protein 1 (IGF2BP1, Hs00198023). Fold change was determined through ΔΔCt analysis with RPLP0 as the housekeeping gene. <u>Mouse</u>: insulin-like growth factor 2 mRNA binding protein (Igf2bp1, Mm00501602_m1), interleukin 1 beta (Il1B, Mm00434228_m1), lipocalin 2 (LCN2, Mm01324470_m1), hypoxanthine guanine phosphoribosyl transferase (Hprt, Mm03024075_m1). Fold change was determined through ΔΔCt analysis with Hprt as the housekeeping gene.

### RNA sequencing and data processing

Total RNA from whole-thickness, distal ileum was isolated and quality controlled, as described above. Libraries were prepared using a stranded Total RNA-seq library preparation protocol with ribosomal RNA (rRNA) depletion (Illumina, San Diego, CA). Libraries were sequenced on an Illumina NovaSeq platform, generating 100 base-pair paired-end (100PE) reads.

### Read alignment and quantification

Initial quality assessment of raw FASTQ files was performed using *FastQC* (31). Trimmed reads were aligned to the human reference genome (GRCh38) using STAR (32) with default parameters for stranded paired-end data. Transcript-level quantification was performed using Salmon (33).

### Normalization and differential expression analysis

Downstream analyses were carried out in R using the DESeq2 package (34). Raw counts were normalized, accounting for sex and experiment number as covariates. Principal component analysis (PCA) was used to assess potential batch effects. Several samples collected from a more proximal region of the intestine failed to cluster into NEC or Dam groups, and were discarded from the analysis (*n* = 6). Batch correction was applied using the limma package’s removeBatchEffect function (35) for sex and experiment number. Differential expression analysis was then performed using negative binomial generalized linear models as implemented in DESeq2.

### Data visualization

Normalized expression values were visualized using heatmaps, PCA plots, and volcano plots generated with pheatmap, ggplot2, and EnhancedVolcano.

### Data Availability

### Enrichr pathway analysis

Within each genotype, differentially regulated genes with respect to NEC were compared using ENRICHR (https://maayanlab.cloud/Enrichr/). Transcription pathways were assessed using the TRUSTT 2019 database.

### Western Blot Analysis

RIPA-SDS lysate was loaded with Laemmli buffer (final concentration: 1x) on 4%–12% Bis-Tris mini protein gels (1.5 mm NuPAGE™, Invitrogen, ThermoFisher). Gels were resolved in 1X NuPAGE™ MOPS SDS running buffer (Invitrogen, ThermoFisher) for 120 minutes at 120 V. Proteins were transferred to a 0.45 µm PVDF membrane (Immobilon, Millipore Sigma) via a wet transfer system in 1X NuPAGE™ transfer buffer (Invitrogen, ThermoFisher) at 120 V for 60 min. Membranes were blocked with LI-COR PBS blocking buffer (LI-COR, Lincoln, NE) at room temperature for 60 min. The following primary antibodies were applied for 16 hours at 4 °C: anti-Imp1 (Cell Signaling Technologies, Danvers, MA. CAT: 2852, 1:1000, rabbit, polyclonal), anti-GADPH (Millipore Sigma, CAT: MAB374, 1:10,000, mouse, monoclonal), anti-Spdef (Lifespan Biosciences, Inc., Shirley, MA, CAT: LS-C749124, 1:500, rabbit, polyclonal). LICOR IRDye 700, 800 were used as secondaries with 0.02% SDS according to the manufacturer’s recommendation, and visualized with LICOR Odyssey system (LI-COR). Densitometry was performed using Image Studio Lite v. 5.2 (LI-COR).

### Immunofluorescence Staining

Designated ileal tissue samples (0.5 cm) were fixed in 4% paraformaldehyde (Electron Microscopy Sciences, Hatfield, PA) for 24 hours, washed three times with PBS, and stored in 70% ethanol until paraffin embedding. Sections (5μm thick) were mounted on glass slides. Staining protocol is modified from Digrazia et al (36). Slides were incubated at 60 °C for 10 minutes, cooled to room temperature, and then deparaffinized in Histoclear (three times, 10 minutes each) followed by rehydration through graded ethanol baths. Rehydration was performed sequentially in 100% ethanol (three times), 95% ethanol (two times), and 75% ethanol (once). Antigen retrieval was performed under high pressure (Cuisinart 12-1 Multicooker, Stamford, Connecticut) in citrate buffer (pH 6.0) for 45 minutes. Nonspecific binding was blocked using 5% bovine serum albumin (BSA) (GoldBio, Saint Louis, MO) or 5% fish gelatin with 0.1% Triton-X (Fisher Bioreagents, Waltman, MA) in PBS for 90 minutes at room temperature. Slides were incubated with anti-Muc2, GTX100664, (GeneTex, Irvine, CA) overnight at 4 °C. Following three washes with PBS, sections were incubated with secondary antibodies, Goat anti-Rabbit IgG, A-11008 (Invitrogen, ThermoFisher) at a dilution of 1:200 and or UEA I DyLight 649 (Vector Laboratories, Carlsbad, CA) at 10 µg/ml for 1 hour in a humidified chamber at room temperature. Nuclei were stained with DAPI (1 μg/mL) for 5 minutes. Slides were mounted using Prolong Gold antifade reagent (Invitrogen, Carlsbad, CA) mounting medium.

### Muc2 quantification

Muc2-positive mucus granules were counted per crypt/villus unit using FIJI. Crypt/villus units were only considered for counting if villi architecture was intact and clearly visible for at least 45 µm into the lumen. To maintain consistency, Muc2-positive granules were only counted within the first 45 µm of the villus measured from crypt base. Each animal represents the average of 6-40 units counted per animal across 3-4 fields of view.

### UEA1 quantification

Acquired images were segmented to mucus granule ROIs using ilastik (version 1.4.0) supervised pixel classification by entire field of view (FOV). Average ROI pixel intensities were measured using FIJI. Each animal’s final intensity represents the mean of 2-3 FOVs.

### Phloxine-Tartrazine/Alcian Blue staining

Phloxine-Tartrazine/Alcian Blue staining of paraffin sections of mouse small intestine of 5 µm thickness was performed as described by Lendrum (Lendrum, 1947). Briefly, deparaffinized sections were exposed to 3% (v/v) acetic acid at room temperature for 3 min and then to 1% (w/v) Alcian Blue (8GX, A3157, Sigma-Aldrich, St. Louis, MO, USA) for 30 min. Subsequently, the sections were given a 10 s exposure of 3% (v/v) acetic acid and a 10 min wash of running water. For nuclear staining, the sections were then stained with 1x Harris’s Hematoxylin (26041-05, Electron Microscopy Sciences, Hatfield, PA, USA) for 20 s and then after a 3 min wash of running water they were stained with 0.5% (w/v) Phloxine B (41787-0250, Acros Organics, Geel, Belgium), 0.5% (w/v) calcium chloride solution for 20 min. After a 1 min wash of running water, sections were differentiated in their staining for image development with exposure to a saturated solution of 1.5% (w/v) Tartrazine (A17682.14, Thermo Scientific, Waltham, MA, USA) in Cellosolve solution (ethylene glycol monoethyl ether; 2-ethoxyethanol) (26106-05, Electron Microscopy Sciences, Hatfield, PA, USA) for 3-20 min. Stained sections were then exposed to 100% (v/v) ethanol for 3 min, incubated in 100% (v/v) xylene for 5 min and mounted with Cytoseal XYL xylene mounting medium (8312-4, Epredia, Kalamazoo, MI, USA). Stained sections were then imaged by brightfield microscopy using an Olympus BX53 microscope (Center Valley, PA, USA). At least 10 well-sectioned crypts were used for counting Phloxine+ Paneth cells, Alcian Blue+ goblet cells, and Phloxine-Tartrazine/Alcian Blue+ (PTAB+) hybrid Paneth/goblet cells.

### Histological damage scoring

Hematoxylin and eosin-stained sections were scored by OHSU pathologist Melanie Hakar, DO, who was blinded to the genotype and treatment of the animals. Damage scores were assigned as follows: 0 = no abnormality; 1 = mild vacuolization; 2 = full-thickness vacuolization; 3 = vacuolization with basal detachment of epithelium. n = 12-16 mice per group.

### Microscopy

Widefield images were taken using a Keyence BZ-X800 (Keyence, Itasca, IL, USA) with a 20X 0.75 NA objective. Fluorescence intensity was quantified using ImageJ.

### RNA-binding protein database

In silico analysis of Imp1 binding to validated and experimental targets was conducted using the RNA-binding protein database (http://rbpdb.ccbr.utoronto.ca/index.php) v1.3 release 28.09.2012. Mouse 3’ UTR and coding (CDS) sequences from Ensembl were used to query predicted Imp1 binding. Known targets *Actb* (37) and *Itgb5* (14) (3’ UTR binders) and *Kras* (38, 39) (CDS) were used for comparison.

### Statistical analyses

All statistical analyses were performed using Prism Version 10.6.1. A paired t-test was used to test the effect of inflammation on IMP1 expression in the human enteroid samples. The D’Agostino & Pearson test was used to test for normality. An unpaired t-test was used to test for single variable effects in normally distributed data. A Mann-Whitney test was used to test for single variable effects in non-normally distributed data. A two-way ANOVA with Turkey’s multiple comparisons test was used to examine the effect of two variables (genotype and NEC) in the mouse tissue samples.

### NEC-associated enteric bacteria upregulate *IMP1* expression in neonatal human enteroids

Using our NEC-in-dish ex vivo model of inflammation (in preparation), we stimulated apical-out neonatal human enteroids with a UV-killed, NEC-associated enteric bacterial culture isolated from a patient with NEC *totalis* (29). UV-treatment preserves inflammatory pathway stimulation, upregulating *IL8* (Figure 2A), without risk of cross-culture contamination. In this inflammatory setting, *IMP1* mRNA was increased 120% relative to PBS-treated controls (n=4; p=0.03, by paired t-test) (Figure 2B). This suggests that in early neonatal intestine, IMP1 is upregulated in response to inflammatory damage. To delineate the implications and downstream molecular consequences of IMP1 upregulation during NEC, we turned to our tissue-specific, genetic mouse models.

**Figure 2.**
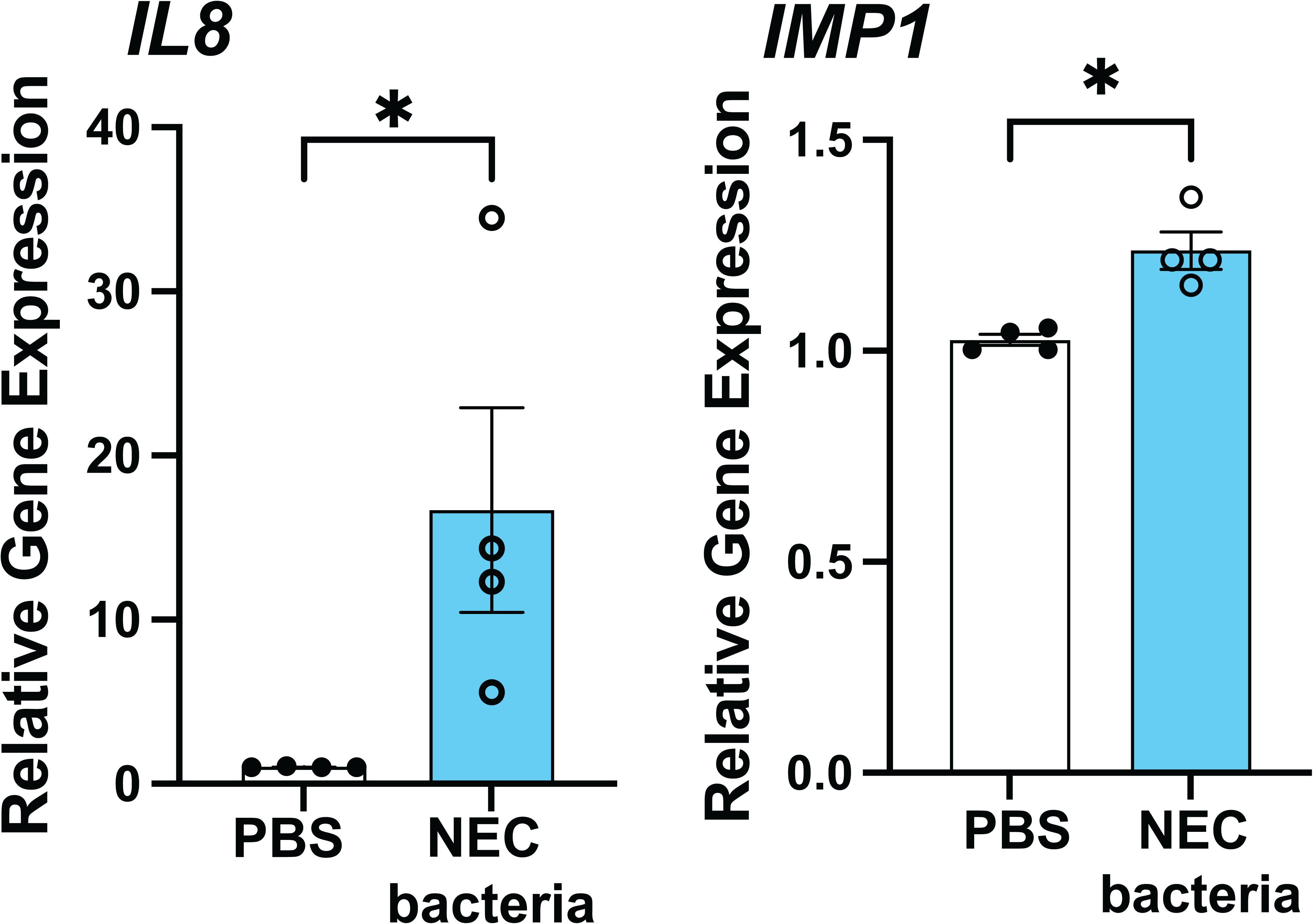
Inflammatory bacteria induce IMP1 expression in neonatal human enteroids. Neonatal human enteroids were flipped apical-out via suspension culture and then treated for 8 hours with a UV-killed NEC-associated enteric bacteria isolated from a patient with NEC *totalis* (29) to induce an inflammatory response or phosphate-buffered saline (PBS) as a control. A) Levels of inflammatory cytokine *IL8* and B) *IMP1* RNA were assessed by quantitative reverse transcription polymerase chain reaction (qRT-PCR) and normalized to housekeeping gene *RPLP0*. Each data point represents one participant sample, n=4. *IL8* data were not normally distributed and were analyzed by a lognormal Welch’s t-test. *IMP1* data were normally distributed and were analyzed by a paired t-test. Error bars represent the SEM and * P <0.05 was considered statistically significant.

### Intestinal epithelial cell (IEC)-specific Imp1 knockout and overexpression mice exhibit expected changes in Imp1 expression

We generated intestinal epithelial-specific Imp1 overexpressing mice (Imp1^IEC-OE^) by crossing mice carrying a STOP sequence flanked by loxP sites upstream of the Imp1 coding sequence (9) to constitutive Villin-Cre. Villin-Cre is active as early as embryonic day 9 (40), therefore, Imp1 is altered before endogenous expression begins at embryonic day 10.5 (1). In the presence of Cre, the STOP sequence is removed, resulting in a 16.4-fold increase in Imp1 mRNA (Figure 3A) and a 4.3-fold increase in Imp1 protein in the intestinal epithelial cells (Figure 3B, *P*<0.05 by unpaired Student’s t-test vs wild-type, littermate controls). We also generated intestinal epithelial cell-specific Imp1 knockout mice (Imp1^ΔIEC^) using mice originally described in Hamilton et al (11) These mice carry homozygous Imp1 alleles where exons 5-6 are flanked by loxP sites (fl/fl). Crossing these Imp1^fl/fl^ mice to constitutive Villin-Cre resulted in a 3.8-fold decrease in Imp1 mRNA and a 30-fold decrease in Imp1 protein in the intestinal epithelial cells (Supplemental Figure 1). The presence of faint bands in the Imp1^ΔIEC^ may be due to the presence of non-epithelial cell types with intact Imp1 expression.

**Figure 3.**
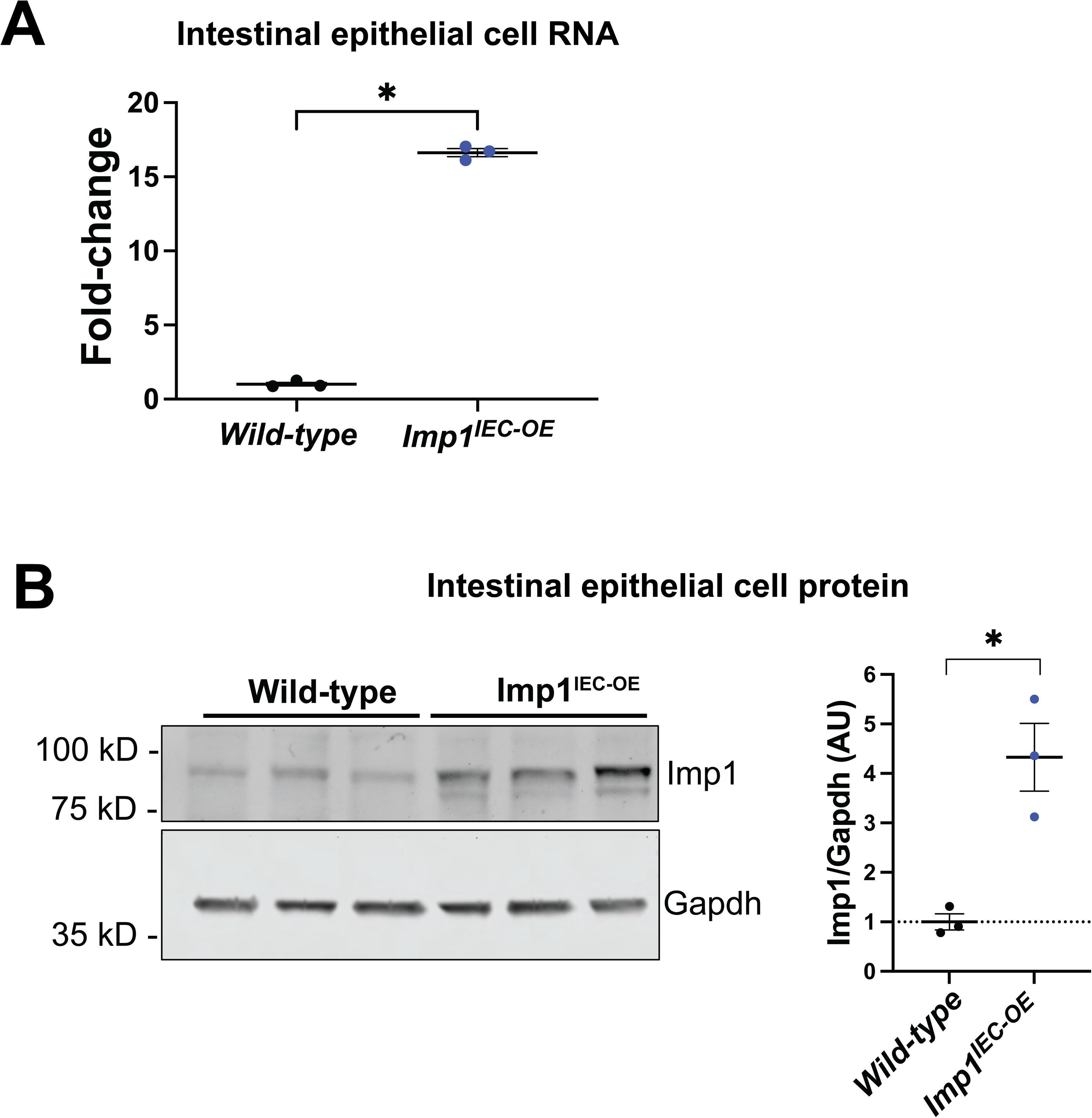
Mouse models of intestinal epithelial-specific Imp1 overexpression or deletion display the expected expression pattern. Intestinal epithelial cells (IECs) were isolated from mice of the following genotypes: VillinCre;lsl-FLAG-Imp1 (Imp1^IEC-OE^), overexpressing Imp1 only in intestinal epithelial cells; and littermate controls with unmodified Imp1 expression. The genotypes of wild-type animals were VillinCre or lsl-FLAG-Imp1. A) Levels of Imp1 RNA were assessed by quantitative reverse transcription polymerase chain reaction (qRT-PCR) in IEC preparations from Imp1^IEC-OE^ and wild-type littermate controls. B) Levels of Imp1 protein in isolated IECs from Imp1^IEC-OE^ and wild-type littermate controls were measured by western blotting. Protein levels were quantified using the area under the curve determined by Image Studio Lite version 5.2 and normalized to the loading control (Gapdh). Each data point represents one animal, n=3 per group. Data were analyzed by Student’s t-test. Error bars represent the SEM and * P <0.05 was considered statistically significant.

### NEC Mouse Model Recapitulates NEC-like Intestinal Injury

To examine the role of Imp1 in the intestinal epithelium during early damage, we used a validated mouse model of NEC-like intestinal injury (30, 41). NEC was induced using a combination of stress, formula feeding, bacterial exposure, and hypoxia beginning at postnatal day 3. Formula-fed mice developed hallmark features of human NEC, including pneumatosis intestinalis, which was absent in dam-fed controls (Figure 4A) and significantly elevated intestinal expression of the inflammatory cytokines *Il1b* and *Lcn2* (Figure 4B and Supplemental Figure 2). Notably, the phenotype of Imp1^ΔIEC^ mice with NEC was similar to that of wild-type animals; therefore, we have placed the Imp1^ΔIEC^ animal data in the supplement for comparison, but limited the main text of the manuscript to a detailed analysis of the more striking Imp1^IEC-OE^ phenotype. The histograms in the supplement include the data from the wild-type and Imp1^IEC-OE^ groups shown in the main text for ease of comparison.

**Figure 4.**
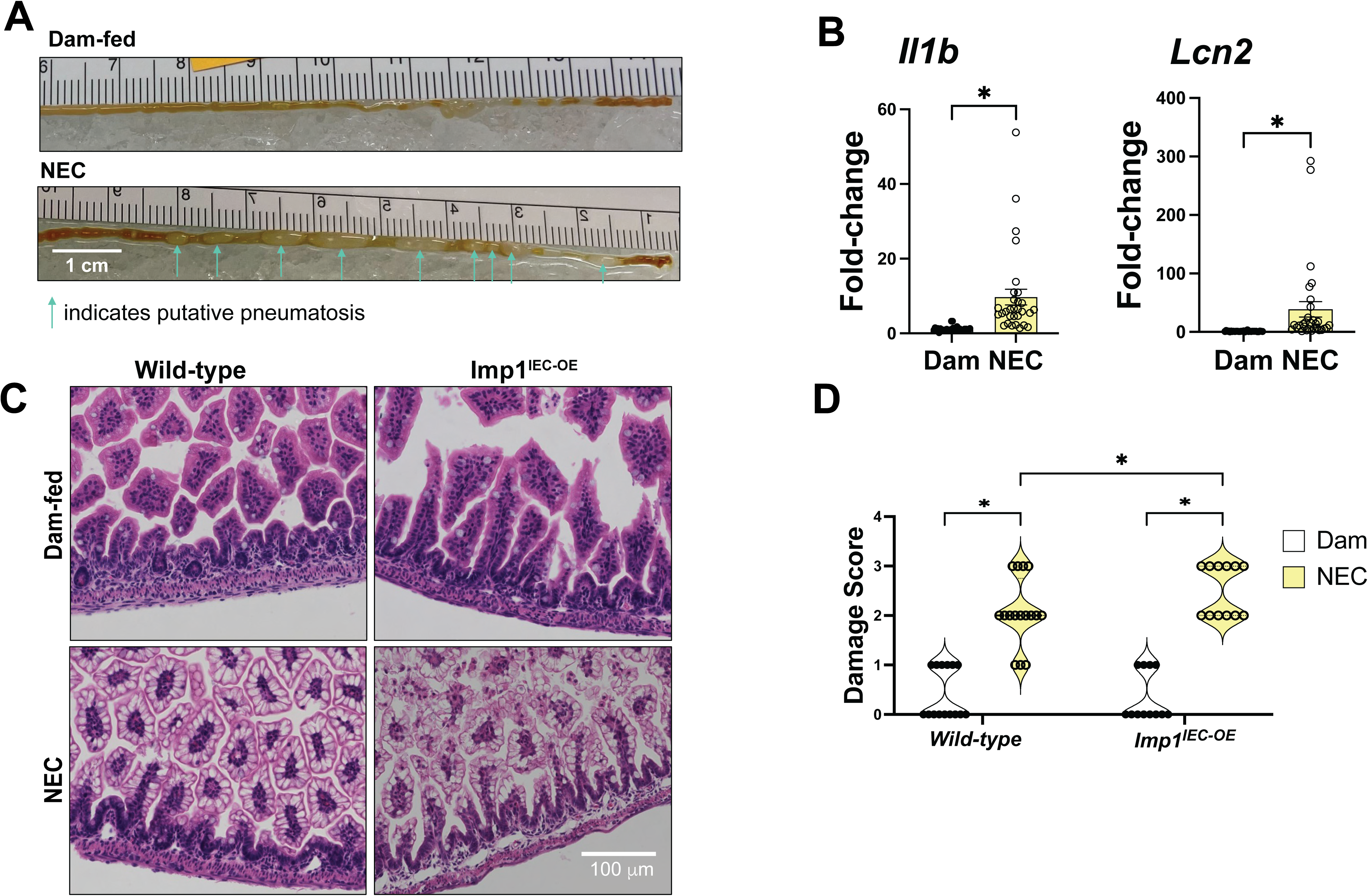
Mouse model of NEC induces inflammatory cytokine gene expression and visible intestinal damage. A) Gross images of intestine from dam-fed and NEC mice. Putative pneumatosis is indicated by arrows. B) Cytokine gene expression for *Il1b* and *Lcn2* was assessed in whole-thickness intestine using qRT-PCR in Imp1^IEC-OE^ versus wild-type dam-fed and NEC mice. Each data point represents one animal, n=30-31 mice per group. The data are not normally distributed and were therefore analyzed using the Mann-Whitney test. Error bars represent the SEM, * P <0.05. C) NEC mice exhibit epithelial vacuolization and damage assessed by hemotoxylin & eosin (H&E) staining and scored D) by a pathologist blinded to animal genotype and treatment. Scores were assigned as follows: 0 = no abnormality; 1 = mild vacuolization; 2 = full-thickness vacuolization; 3 = vacuolization with basal detachment of epithelium. Each dot represents one animal, n = 12-16 mice per group. Data were analyzed by a 2-way ANOVA with Tukey’s test for multiple comparisons. Error bars represent the SEM and * P <0.05 was considered statistically significant.

### IEC-Imp1 overexpressing mice exhibit worse histological damage during NEC

We used hemotoxylin and eosin staining to evaluate the intestinal morphology and compare the histological damage between wild-type and Imp1^IEC-OE^ mice during NEC. Histological damage scoring was performed by a pathologist who was blinded to the genotype and condition of the mice. Scores revealed that Imp1^IEC-OE^ mice exhibit more severe histological damage during NEC, as defined by vacuolization with detachment from the basement membrane (Figure 4C-D).

### IEC-Imp1 overexpression promotes Spdef transcriptional activity in response to NEC

We performed RNA sequencing analysis on whole-thickness ileal tissue isolated from wild-type and Imp1^IEC-OE^ mice after NEC induction and dam-fed littermate controls. Principal component analysis (PCA) revealed distinct clustering based on experimental cohort (NEC mice vs. dam-fed controls) across all genotypes (Figure 5A; Supplemental Figure 3A). Imp1 expression differences were preserved across treatment groups (Supplemental Figure 3B), with Imp1 remaining significantly elevated in Imp1^IEC-OE^ mice regardless of treatment (Figure 5B). Globally NEC induction up-regulated secretory gene expression within the ileum (Supplemental Figure 4). We then used Enrichr to investigate global changes in transcription factor pathways in response to NEC within each genotype group. Wild-type and Imp1^IEC-OE^ mice up-regulated the Rela pathway in response to NEC. Uniquely, Imp1^IEC-OE^ mice exhibited transcriptional signatures linked to up-regulation of caudal type homeobox 2 (Cdx2) and SAM Pointed Domain Containing ETS Transcription Factor (Spdef) (Figure 5C). Both Cdx2 (42) and Spdef (43–48) are critical for IEC differentiation. Spdef is required for goblet cell differentiation and is responsible for mature goblet cell function (43–49). *Spdef* mRNA is significantly up-regulated in both genotypes in response to NEC (124.5 ± 10.9 vs 241.0 ± 30.0 [WT]; 103.0 ± 8.2 vs 338.0 ± 49.0 [Imp1^IEC-OE^]; however, the level of expression is significantly higher (p=0.03 by 2-way ANOVA vs WT NEC) in Imp1^IEC-OE^ during NEC (Figure 5D). This translates to a 1.5-fold increase in Spdef protein in the Imp1^IEC-OE^ during NEC relative to wild-type NEC mice (Figure 5E). Spdef co-regulates IEC fate in conjunction with atonal homolog 1 (Atoh1) (50). *Atoh1* was also up-regulated in response to NEC, but not differentially based on genotype (Figure 5F). Importantly, known Spdef transcriptional targets, *Gcnt3*, *Agr2,* and *Foxa3* (44, 49) are significantly up-regulated in Imp1^IEC-OE^ above levels observed in WT mice during NEC (Figure 5G). Collectively, these results led us to ask whether *Spdef* is a direct target of Imp1.

**Figure 5.**
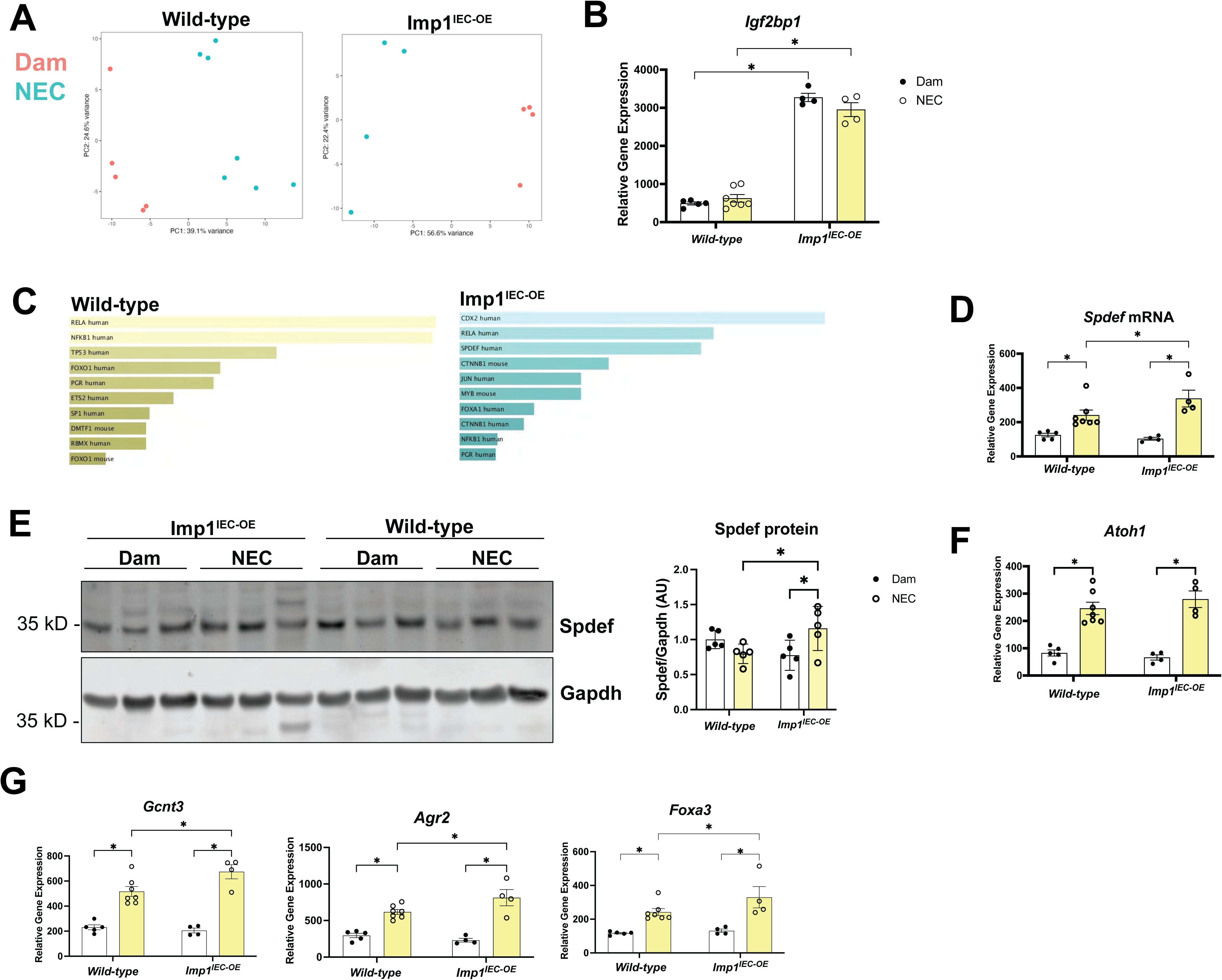
Intestinal epithelial cell (IEC) Imp1 up-regulates Spdef and Spdef transcriptional program in response to NEC injury. Total RNA was isolated from the ileum (whole-thickness) of dam-fed and NEC mice and used for RNA sequencing and analysis. A) Principal component analysis (PCA) plots assess data stratification based on dam-fed vs. NEC induction within each genotype group. Variance stabilizing transformation (VST) normalized counts for all (n = 20) wild-type and NEC-induced samples accounting for sex and experiment number as covariates. The samples are colored in blue for NEC and red for Dam, respectively. The percent of variation explained is indicated for principal component 1 along the x axis and principal component 2 along the y axis. B) Imp1 gene expression was confirmed across genotypes and treatment groups. C) ENRICHR analysis assessed differential transcription factor pathway activity between dam-fed and NEC conditions for each genotype. Data shown are from the TRUSTT 2019 database and are sorted by p-value. D) *Spdef* and E) *Atoh1* gene expression were compared between dam-fed and NEC wild-type and Imp1^IEC-OE^ mice using RNA sequencing. F) Protein was isolated from the ileum (whole-thickness) of dam-fed and NEC mice of each genotype and analyzed for Spdef using western blotting. Protein levels were quantified using the area under the curve determined by Image Studio Lite version 5.2 and normalized to the loading control (Gapdh). G) Gene expression for Spdef transcriptional targets *Gcnt3*, *Agr2*, and *Foxa3* was compared between dam-fed and NEC wild-type and Imp1^IEC-OE^ mice using RNA sequencing. All data were analyzed by a two-way ANOVA with Tukey’s test for multiple comparisons to determine the effect of NEC and Imp1 on the parameters measured. Each data point represents one animal, n=4-7 per group. Error bars represent the SEM and * P <0.05 was considered statistically significant.

### Imp1 up-regulates Spdef and is predicted to bind to the *Spdef* coding sequence

We utilized the RNA-binding protein database (51) to query if Imp1 is predicted to bind *Spdef* mRNA. We focused on the coding sequence and 3’ untranslated region, which is the most common binding location for Imp1 (14, 15). We found that Imp1 is predicted to bind *Spdef* coding sequence with a binding score of 8.83, which is higher than the score for a known Imp1 target, *Kras* (38, 39), which had a binding score of 7.06 (Table 2). Predicted binding values for the 3’ UTR and known 3’ UTR targets *Actb* (37, 52) and *Itgb5* (14) are shown for comparison. Taken together these data indicate that Imp1 may bind to and stabilize Spdef mRNA during early neonatal intestinal inflammation. Given the critical role of Spdef in goblet cell differentiation and differentiated function, we next asked if goblet cell number was changed with respect to Imp1 expression during NEC.

**Table 2.** RNA-binding protein database predicted Imp1 binding to Spdef relative to known Imp1 targets.

| Gene | CDS |  |  | 3' UTR |  |  | Source |
| --- | --- | --- | --- | --- | --- | --- | --- |
|  | Score | Relative Score | Potential Targets | Score | Relative Score | Potential Targets |  |
| **Actb** | 8.65 | 49% | 17 | 7.35 | 41% | 12 | Nicastro et al., 2017, Cell Rep; Huttelmaier et al., 2005, Nature |
| **Itgb5** | 5.7 | 32% | 33 | 4.63 | 26% | 10 | Conway et al, 2016. Cell Rep. |
| **Kras** | 7.06 | 40% | 7 | 5.01 | 28% | 6 | Mackedenski et al., 2018, Biochem J |
| **Spdef** | 8.83 | 50% | 18 | 4.20 | 23% | 3 | Experimental |
| **Qsox1** | 5.74 | 32% | 25 | 10.95 | 62% | 3 | Experimental |

\*\*Experimentally validated targets are shown in blue.

### Crypt-base goblet cells expand during NEC, irrespective of genotype

We analyzed goblet cells using Muc2 immunofluorescent staining and quantified goblet cells within or near the intestinal crypts. We found that NEC induced goblet expansion in the crypt; however, the magnitude of goblet cell expansion did not differ between the wild-type and Imp1^IEC-OE^ mice during NEC. Notably, there was a significant increase in crypt-base goblet cells in the Imp1^IEC-OE^ after NEC, but this increase was driven by a slight, non-significant decrease in crypt-based goblet cells in the homeostatic condition (Figure 6A-B). For a confirmatory analysis, we used phloxine tartrazine-alcian blue (PT-AB) staining to identify Paneth cells stained with phloxine tartrazine, goblet cell acidic mucins stained with alcian blue, and intermediate or hybrid cells that label with both stains (Figure 6C). We quantified each cell type and found that alcian blue-positive goblet cells were also expanded during NEC and that the significant increase seen in the Imp1^IEC-OE^ mice was driven by a slight decrease in alcian blue-positive cells at homeostasis (Figure 6D). Paneth cells also expanded in the NEC condition, but the increase was only significant in the wild-type group (Supplemental Figure 5). The number of hybrid cells was small and did not differ by genotype or inflammatory state (Supplemental Figure 5). These data align with RNA sequencing analysis of common goblet cell markers, *Clca1*, *Tff3*, *Muc2*, *Fcgbp*, and *Zg16*, which are increased (*P*<0.05) during NEC, but do not differ between genotypes (Figure 6E). Goblet cells differentiate from actively dividing stem cells (53) and interestingly, active stem cell marker, Lgr5, is upregulated in Imp1^IEC-OE^ mice at homeostasis (Figure 6F). These data suggest that the increased expression of goblet cell-specific Spdef targets relative to wild-type is not mediated by an increase in goblet cell number. Given the role of Spdef in mediating differentiated goblet cell function, we then asked if genes involved in mature goblet cell function differed between wild-type and Imp1^IEC-OE^ mice during NEC.

**Figure 6.**
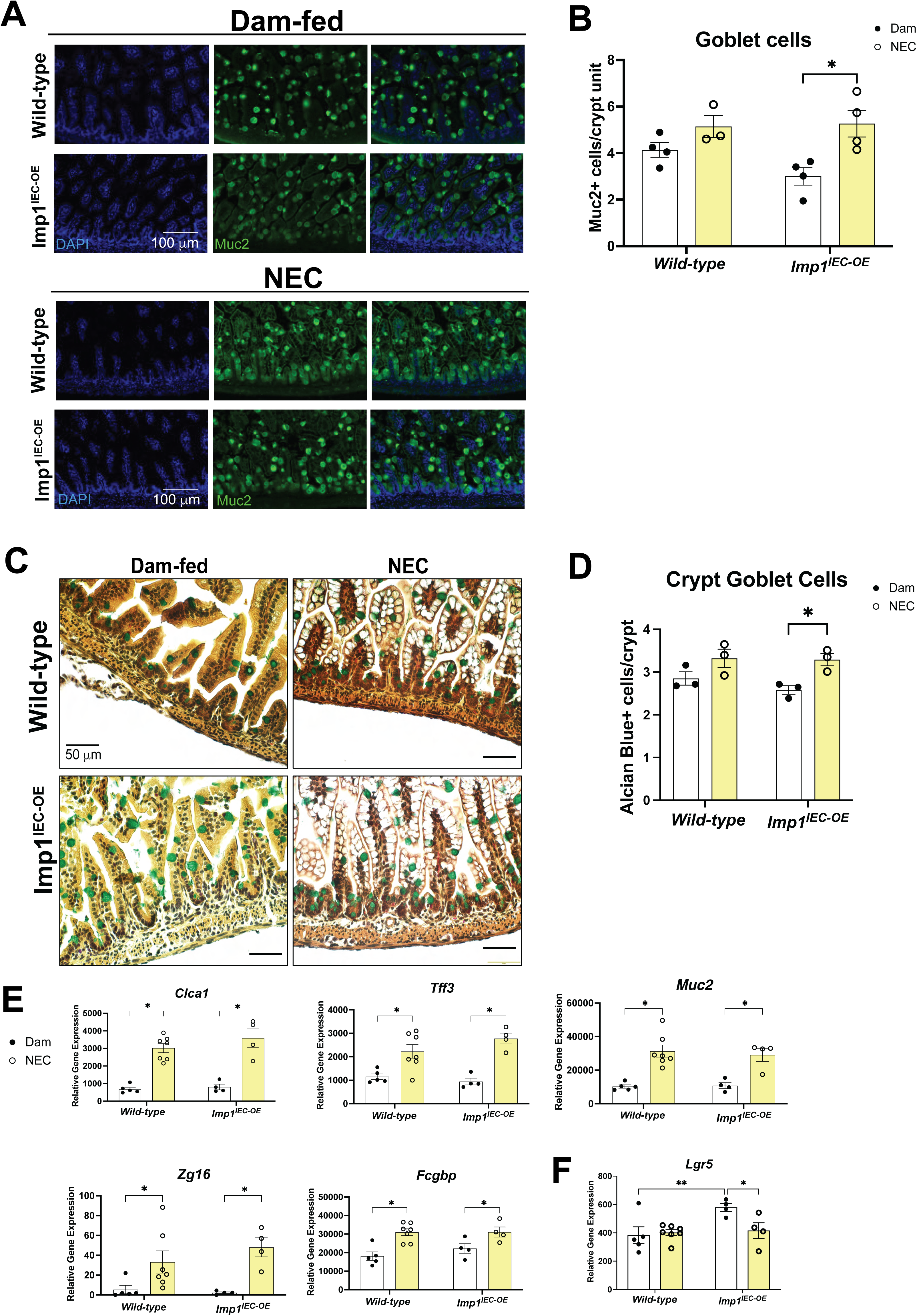
NEC promotes crypt-base goblet cell expansion and associated gene expression. A) Immunofluorescence staining for Muc2 in the ileum of wild-type and Imp1^IEC-OE^ in dam-fed and NEC groups. B) Crypt-associated Muc2-positive cells were quantified by an observer blinded to the genotype and treatment, n=3-4 animals. C) Phloxine-tartrazine alcian blue staining was used to assess goblet cells (alcian-blue positive) in the ileum of wild-type and Imp1^IEC-OE^ in dam-fed and NEC groups. D) Alcian-blue positive cells within the crypts were quantified by an observer blinded to the genotype and treatment, n=3 animals. E) Total RNA was isolated from the ileum (whole-thickness) of dam-fed and NEC mice for RNA sequencing. Analyses assessed goblet cell markers in wild-type and Imp1^IEC-OE^ mice from dam-fed and NEC groups. n = 4-7. For all data shown, error bars represent SEM. P-values were determined by 2-way ANOVA with Tukey’s test for multiple comparisons; * P < 0.05 was considered statistically significant.

### Imp1 promotes goblet cell maturation and mucus fucosylation during NEC

Using the RNA sequencing dataset, we compared the expression of genes involved in differentiated goblet cell function, mucus glycosylation, and mucus production/secretion between wild-type and Imp1^IEC-OE^ mice during NEC. Notably, all were significantly upregulated in Imp1^IEC-OE^ mice during NEC (Figures 6). *Spink4, Klk1, Tspg1, Ccl6, Defa24, Creb3l4* are Spdef-responsive genes linked to differentiated goblet cell function (45), further supporting a role of Imp1 in regulating Spdef during NEC (Figure 7A). The production of heavily glycosylated mucus is a chief function of goblet cells also closely linked with Spdef (44–46). Consistently, we observe increased expression of genes involved in mucus glycosylation, including *Agr2* and *Gcnt3* (Figure 4G) and *Fut2, B3gnt7,* and *Qsox1* (Figure 7B). Imp1 regulation of mature goblet cell functions may extend beyond Spdef, as Imp1 is also predicted to bind *Qsox1* mRNA (Figure 7C and Table 2), based on published enhanced crosslinking immunoprecipitation (eCLIP) data (14) and the RNA binding protein database. Mining eCLIP data (14) also suggests that IMP1 may bind *B3GNT7* and *FUT2*. These gene expression differences in glycosylation mediators led us to examine the effect of Imp1 on mucus glycosylation during NEC using lectin staining for ulex europaeus agglutinin-1 (UEA1), which measures the presence of fucose added by Fut2. UEA1 staining results show increased levels of UEA1 in Imp1^IEC-OE^ mice during NEC (957.1 ± 181.6 vs 590.3 ± 49.7, dam-fed Imp1^IEC-OE^ (*P* < 0.05) and 686.5 ± 31.42 wild-type NEC mice (*P* = 0.07) (Figure 7D & E).

**Figure 7.**
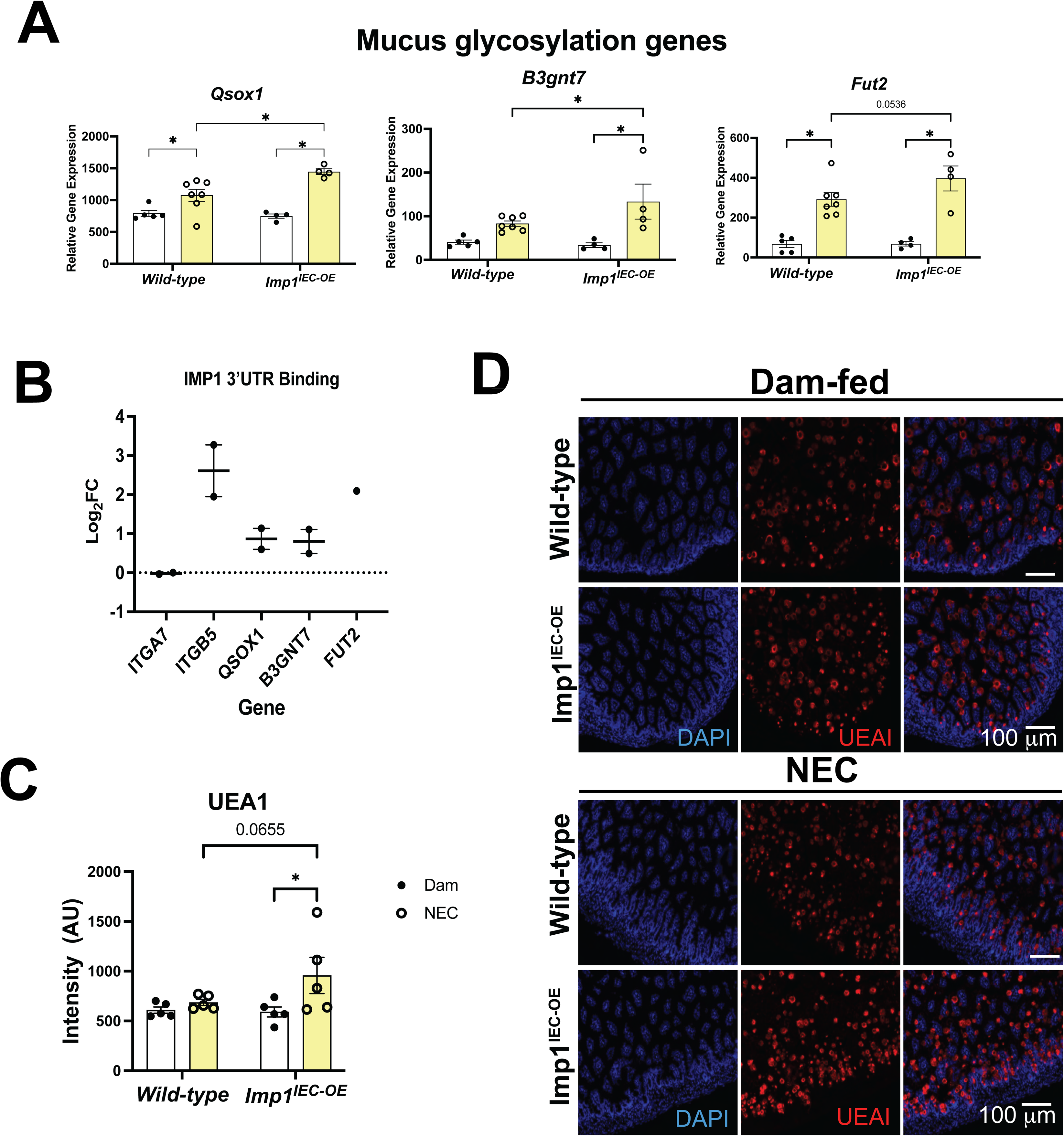
Imp1 promotes fucosylated mucus production during NEC. Total RNA was isolated from the ileum (whole-thickness) of dam-fed and NEC mice for RNA sequencing. Analyses assessed A) genes associated with Spdef (45) or B) additional genes with roles in goblet cell mucus glycosylation in wild-type and Imp1^IEC-OE^ mice from dam-fed and NEC groups. n = 4-7. C) An IMP1 enhanced crosslinking immunoprecipitation (eCLIP) dataset from induced pluripotent stem cells was mined and relative 3’ untranslated region (UTR) binding scores are shown. ITGA7 and ITGB5 are known negative (7) and positive (14) controls shown for reference. D) UEA1-positive cells were quantified to determine the effect of genotype and treatment on mucus fucosylation. n=5 mice per group. E) Representative UEA1 immunostaining compares the amount of fucosylated mucus (mediated by Fut2) between wild-type and Imp1^IEC-OE^ mice in the dam-fed and NEC groups. For all data shown, each data point represents one mouse and error bars represent SEM. P-values were determined by 2-way ANOVA with Tukey’s test for multiple comparisons; * P<0.05 was considered statistically significant.

Our findings demonstrate that IMP1 expression is enhanced in human neonatal enteroids exposed to inflammatory, NEC-associated enteric bacteria. In a mouse model of NEC, Imp1 overexpression promotes goblet cell maturation and enhances mucus fucosylation, while exacerbating histological damage during intestinal inflammation. The data suggest that this goblet cell transcriptional program may be mediated via Imp1 stabilization of Spdef mRNA. Additional experiments are required to determine whether Imp1 upregulation is ultimately beneficial or detrimental, as well as to demonstrate that *Spdef* and mucus glycosylation mediators are targets of Imp1 during intestinal inflammation.

## DISCUSSION

This study investigated the importance of the RNA-binding protein Imp1 in mediating neonatal intestinal inflammatory responses using human enteroids and a mouse model of NEC. Our results indicate that IMP1 is upregulated in neonatal, human enteroids in response to inflammatory bacteria. Using animal models with IEC Imp1 loss and overexpression we demonstrate that the upregulation of Imp1 results in worse IEC damage during NEC, but promotes a Spdef transcriptional program, goblet cell maturation, and enhanced fucosylated mucus during NEC inflammation. We speculate that these transcriptional and translational changes may be driven through direct Imp1 binding to *Spdef* itself or the mRNAs of other regulators of goblet cell mucus glycosylation. These findings establish an intriguing paradox where elevated Imp1 expression results in a more damaged epithelium while also promoting goblet cell maturation and mucus fucosylation, adaptations that are generally considered protective in the setting of inflammation (26, 27).

Imp1 is thought to play important roles in intestinal development (1) and is differentially regulated in response to inflammation across pediatric and adult inflammatory bowel disease (IBD) (7). This prompted us to examine its role in NEC, which occurs during key intestinal development milestones and is highly inflammatory. Pediatric IBD patients have lower IMP1 levels than uninflamed controls, while adult IBD patients upregulate IMP1 (7). The precise role of IMP1 during intestinal damage is multifaceted. One study using an inducible Villin-Cre system shows that Imp1 loss worsens IBD by disrupting barrier function via occludin downregulation (12) In contrast, several studies using constitutive, tissue-specific Imp1 loss in the intestinal and colonic epithelium demonstrate that the presence of Imp1 worsens damage recovery by blunting autophagy-driven regeneration (7) and facultative stem cell expansion (54). Notably, our NEC model is focused entirely on the damage response, as recovery and repair experiments are inherently challenging. In our hands, outcomes for mice with wild-type Imp1 or Imp1 loss are similar, while Imp1^IEC-OE^ mice exhibit worse intestinal damage. Exacerbated damage in the Imp1^IEC-OE^ mice may be due to changes in autophagy regulation or in IEC barrier function, hypotheses which we are currently investigating. Alternatively, elevated damage in the Imp1^IEC-OE^ mice could be due to changes in the intestinal stem cell (ISC) compartment during homeostasis. At baseline, we observe an increase in *Lgr5* mRNA, indicating the potential for more actively dividing stem cells, which are highly susceptible to damage (55). Recent work also shows that Lgr5-ISCs can differentiate directly into secretory lineages (such as goblet cells) (53), which may further support a role for upregulating *Spdef* to promote goblet cell maturation.

Under homeostatic, uninflamed conditions, Imp1 is dispensable for early goblet cell differentiation or mature gene expression, consistent with published studies in adult mice (8). Similarly, elevated Imp1 levels have no measurable effect on goblet cell development or gene expression in the unperturbed state, as shown previously in adult mice (9). However, after NEC induction, mice with elevated intestinal epithelial Imp1 expression exhibit specific up-regulation of genes linked to goblet cell maturation, including genes related to mucus O-glycosylation (*Fut2, Gcnt3*, *B3gnt7, Qsox1*), genes involved in mucus protein processing (45, 47) (*Agr2, Klk1*), mucus secretion (56, 57) (*Creb3l4*), and other mature goblet cell functions (45, 47, 58, 59) (*Ccl6*, *Spink4*, *Tpsg1, Foxa3*). Notably, not all goblet cell-specific genes are upregulated in this manner. Genes such as *Clca2, Tff3*, *Fcgbp*, *Muc2,*and *Zg16* are expressed at similar levels in wild-type and Imp1^IEC-OE,^ indicating that the changes are not likely the byproduct of a loss of non-goblet cell mRNAs in the RNA sequencing analysis. In fact, most of the significantly upregulated genes are known or associated Spdef transcriptional targets (43–47), strongly suggesting that the enhanced goblet cell gene expression changes in the Imp1^IEC-OE^ mice during NEC may be largely Spdef-mediated.

*In silico* analyses suggest that Imp1 may directly bind Spdef mRNA and possibly the transcripts of mucus glycosylation mediators. Predictive binding to the Spdef mRNA coding sequence indicates that Imp1 is a putative Spdef binder, with a higher binding score than the known target Kras (39). Imp1 is also predicted to bind Qsox1, an enzyme responsible for Golgi glycosyltransferases activation (60). The published induced pluripotent stem cell eCLIP dataset from Conway et al (14), further supports this, indicating that IMP1 may bind *QSOX1,* as well as the transcripts for B3GNT7 and FUT2. If this predicted binding holds up to further experimental testing, this would be a novel finding indicating a role for Imp1 in intestinal mucus regulation in response to inflammatory insult.

NEC is known to disrupt the intestinal barrier, leading to decreased barrier protein gene expression and mislocalization (61, 62). Moreover, deficiencies in the mucus barrier promote infection and susceptibility to more severe inflammation relative to mice with an intact mucus barrier (23, 24). Goblet cells play a crucial role in maintaining the intestinal epithelial barrier, limiting inflammation, and regulating the microbiome (22). Whether our observed changes in goblet cell gene expression and mucus glycosylation are sufficient to preserve some level of intestinal barrier function is currently under investigation. Barrier improvements via mucus would be consistent with several studies indicating that preserving goblet cell number via bovine milk exosome treatment (63), administration of acetate-producing bacteria (64), or loss of Tlr4 (65) attenuates damage in NEC mouse models. Additionally, administration of goblet cell products, such as recombinant trefoil factor 3 (66), or human milk components (67), promotes intestinal barrier function, attenuates histological damage, or reduces mucus permeability in a rat model of NEC.

Interestingly, we observe slight goblet cell expansion in the crypt base of our NEC mice regardless of genotype. This is in contrast to a body of literature suggesting that NEC is associated with goblet cell loss (61, 68–72). In rodents, goblet cell expansion and mucus generation occur between p2 and p9 (73). Perhaps the timing of our model (starting at p3) aligns with maximal goblet cell expansion, which is augmented in response to inflammation. Alternatively, we counted goblet cells within the crypt region only, while other studies measured goblet cells in full crypt-villus units (71), villi only (68), total epithelium (71), or by gene expression (70). These differences in quantification may result in our contrasting findings. Additionally, the level of damage in our model may be considered mild relative to other models as the epithelium is not denuded. Mild damage has been linked to goblet cell expansion in NEC (69).

While NEC animal models are a useful tool, they do not fully recapitulate human NEC. Additional studies are needed to determine if altering mucus glycosylation is a tenable target for NEC treatment. Additionally, the level of Imp1 upregulation in our genetic model is high and exceeds the degree of upregulation observed in the human enteroids.

The goal of this work was to use Imp1 manipulation as a tool to shed light on mechanisms of NEC susceptibility or damage protection and to improve our understanding of NEC pathophysiology. Our data reveal Imp1 as a novel regulator of neonatal goblet cell function via the Spdef transcriptional program and mucus glycosylation mediators during NEC. Ongoing studies will examine direct Imp1 binding targets during neonatal inflammation and impacts on barrier function to further define the mechanism, while also clarifying whether Imp1 or the pathways it influences are inherently beneficial or detrimental in NEC.

## ACKNOWLEDGEMENTS

We wish to thank Dr. Brian Scottoline for his edits to the manuscript and for sharing the human tissue samples for enteroid generation and Dr. Misty Good for sharing the NEC-associated enteric bacteria. We wish to thank Zhao Lai and the Next Gen sequencing core at UT Health Sciences Center San Antonio (UTHSCSA) for performing the RNA sequencing. We thank the OHSU Histopathology Shared Resource for embedding, sectioning, and H&E staining. Figure diagrams were created with BioRender.com.

## FUNDING

SFA is supported by the National Institutes of Health (NIH) grant K01DK129401, R03DK142841-01, R01HD109193, and grants from the Gerber Foundation and the Knight Cancer Institute. The work is also supported by the National Institutes of Health (NIH) grant R21HD104922 (PS).

## COMPETING INTERESTS

The authors have no conflicts of interest to declare.

## Supplemental Figures

**Supplemental Figure 1. NEC-induced damage promotes a secretory gene expression profile in the intestinal epithelium.** Total RNA was isolated from the ileum (whole-thickness) of dam-fed and NEC mice for RNA sequencing. Analyses assessed cell type markers for secretory lineages in wild-type, Imp1^ΔIEC^, and Imp1^IEC-OE^ mice from dam-fed and NEC groups. n = 2-7 mice.

**Supplemental Figure 1. Mouse models of intestinal epithelial-specific Imp1 deletion display cell-specific Imp1 loss.** Intestinal epithelial cells (IECs) were isolated from mice of the following genotypes: VillinCre;Imp1^fl/fl^ (Imp1^ΔIEC^), lacking Imp1 only in intestinal epithelial cells and littermate controls with unmodified Imp1 expression. The genotypes of wild-type animals were Imp1^fl/fl^. A) Levels of Imp1 RNA were assessed by quantitative reverse transcription polymerase chain reaction (qRT-PCR) in IEC preparations from Imp1^ΔIEC^ and wild-type littermate controls. B) Levels of Imp1 protein in isolated IECs from Imp1^ΔIEC^ and wild-type littermate controls were measured by western blotting. Protein levels were quantified using the area under the curve determined by Image Studio Lite version 5.2 and normalized to the loading control (Gapdh). Each data point represents one animal, n=3 per group. Data were analyzed by Student’s t-test. Error bars represent the SEM and *p<0.05 was considered statistically significant.

**Supplemental Figure 2. Inflammatory cytokines are up-regulated in NEC model intestines of mice lacking intestinal epithelial cell (IEC) Imp1.** Cytokine gene expression for *Il1β* and *Lcn2* was assessed in whole-thickness intestine using qRT-PCR between dam-fed and NEC mice within the (A) Imp1^ΔIEC^ line. Each data point represents one animal, n=11-12 per group. The data are not normally distributed and were therefore analyzed using the Mann-Whitney test. Error bars represent the SEM, * P <0.05. B) NEC mice exhibit epithelial vacuolization and damage assessed by hemotoxylin & eosin (H&E) staining.

**Supplemental Figure 3. Mice lacking intestinal epithelial cell (IEC) Imp1 exhibit a Spdef transcriptional program similar to that of wild-type animals.** Total RNA was isolated from the ileum (whole-thickness) of dam-fed and NEC mice and used for RNA sequencing and analysis. A) PCA plots assess data stratification based on dam-fed vs. NEC induction for Imp1^ΔIEC^ mice. Variance stabilizing transformation (VST) normalized counts for all (n = 6) wild-type and NEC-induced samples accounting for sex and experiment number as covariates. The samples are colored in blue for NEC and red for Dam, respectively. The percent of variation explained is indicated for principal component 1 along the x axis and principal component 2 along the y axis. B) Imp1 gene expression was confirmed across genotypes and treatment groups. C) ENRICHR analysis assessed differential transcription factor pathway activity between dam-fed and NEC conditions for Imp1^ΔIEC^ mice. Data shown are from the TRUSTT 2019 database and are sorted by p-value. D) *Spdef* and E) *Atoh1* gene expression were compared between dam-fed and NEC wild-type, Imp1^ΔIEC^, and Imp1^IEC-OE^ mice using RNA sequencing. F) Gene expression for Spdef transcriptional targets *Gcnt3*, *Agr2*, and *Foxa3* was compared between dam-fed and NEC wild-type, Imp1^ΔIEC^, and Imp1^IEC-OE^ mice using RNA sequencing. All data were analyzed by a two-way ANOVA with Tukey’s test for multiple comparisons to determine the effect of NEC and Imp1 on the parameters measured. Each data point represents one animal, n=4-7 per group. Error bars represent the SEM and * P <0.05 was considered statistically significant. Note that data for wild-type and Imp1^IEC-OE^ mice are reproduced here to allow for comparison with corresponding Imp1^ΔIEC^ data.

**Supplemental Figure 4. NEC induction up-regulates secretory gene expression across all genotypes when compared to controls.** Three heatmaps comparing the gene expression of known cell-type markers for secretory cells (enteroendocrine, goblet, and Paneth) in dam-fed controls versus NEC samples across the genotypes: wild-type, Imp1^ΔIEC^, and Imp1^IEC-OE^.

**Supplemental Figure 5. NEC induces Paneth cell expansion in wild-type mice.** Phloxine-Tartrazine/Alcian Blue staining was used to assess goblet cells (alcian-blue positive), Paneth cells (phloxine-tartrazine-positive) and hybrid cells (Phloxine-Tartrazine/Alcian Blue dual positive) in the ileal crypts of wild-type and Imp1^IEC-OE^ in dam-fed and NEC groups. Images shown in Figure 6C. A) Phloxine-tartrazine-positive cells (Paneth cells) and B) phloxine-tartrazine alcian blue dual-positive hybrid cells in the crypts were quantified by an observer blinded to the genotype and treatment, n=3 animals.

**Supplemental Figure 6. Mice lacking intestinal epithelial cell (IEC) Imp1 exhibit goblet cell number and gene expression changes similar to those of wild-type animals.** A) Immunofluorescence staining for Muc2 in the ileum of Imp1^ΔIEC^ mice in dam-fed and NEC groups. B) Crypt-associated Muc2-positive cells were quantified by an observer blinded to the genotype and treatment, n=3-4 animals. Total RNA was isolated from the ileum (whole-thickness) of dam-fed and NEC mice for RNA sequencing. Analyses assessed C) goblet cell markers and D) active stem marker *Lgr5* in Imp1^ΔIEC^ mice from dam-fed and NEC groups. n = 2-7. For all data shown, error bars represent SEM. P-values were determined by 2-way ANOVA with Tukey’s test for multiple comparisons; * P <0.05 was considered statistically significant. Note that data for wild-type and Imp1^IEC-OE^ mice are reproduced here to allow for comparison with corresponding Imp1^ΔIEC^ data.

**Supplemental Figure 7. Mice lacking intestinal epithelial cell (IEC) Imp1 exhibit changes in Spdef-associated and mucus glycosylation genes similar to those of wild-type animals.** Total RNA was isolated from the ileum (whole-thickness) of dam-fed and NEC mice for RNA sequencing. Analyses assessed A) genes associated with Spdef (45) or B) additional genes with roles in goblet cell mucus glycosylation in wild-type, Imp1^ΔIEC^, and Imp1^IEC-OE^ mice from dam-fed and NEC groups. n = 2-7. For all data shown, each data point represents one mouse and error bars represent SEM. P-values were determined by 2-way ANOVA with Tukey’s test for multiple comparisons; * P <0.05 was considered statistically significant.

## Notes

### Competing Interest Statement

The authors have declared no competing interest.

