## Supplementary figures and images for "The RNA-binding protein Imp1 promotes a Spdef transcriptional program and mucus fucosylation during necrotizing enterocolitis"

### Supplemental Figure 1

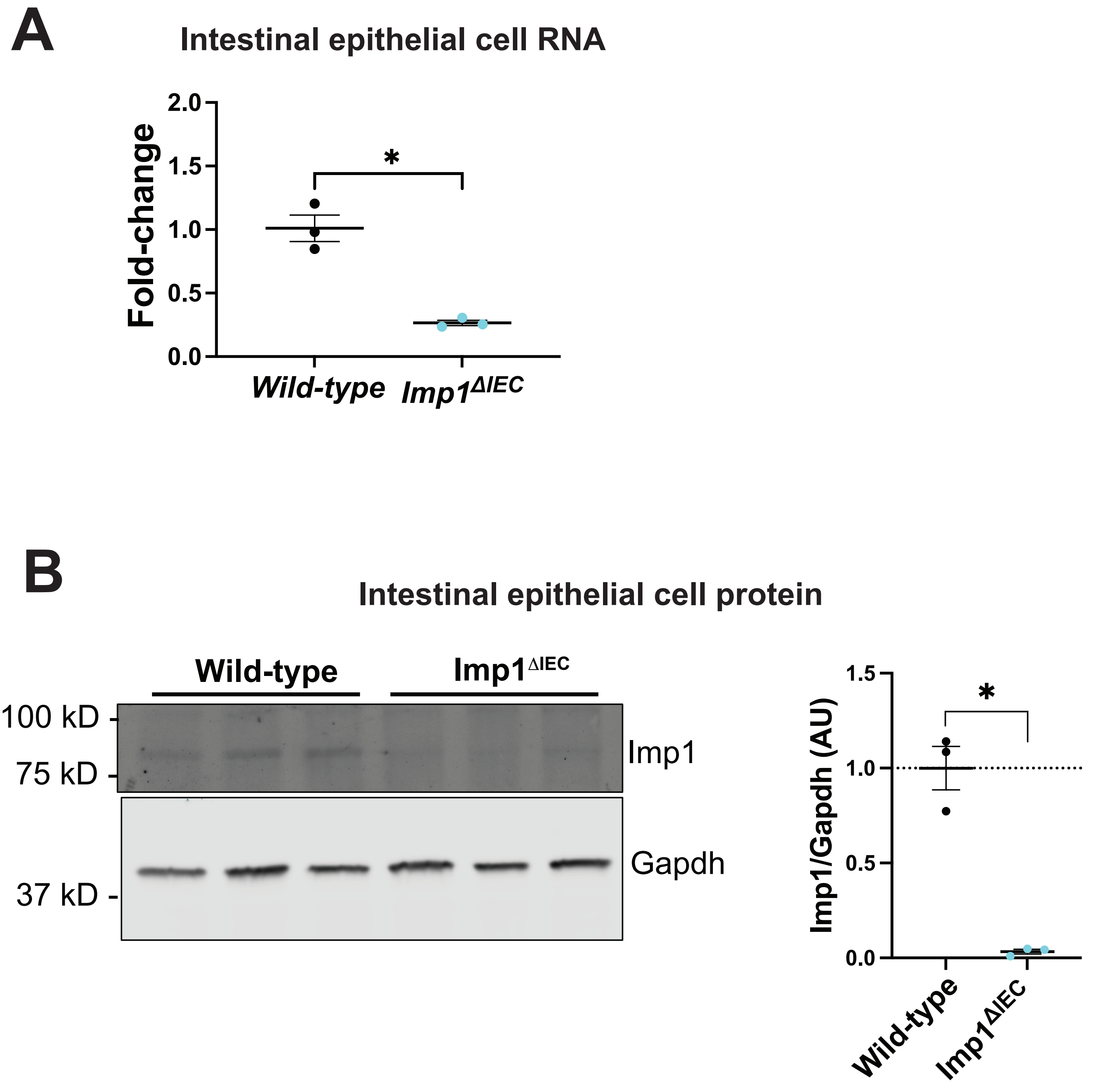


Supplemental Figure 1.

### Supplemental Figure 2

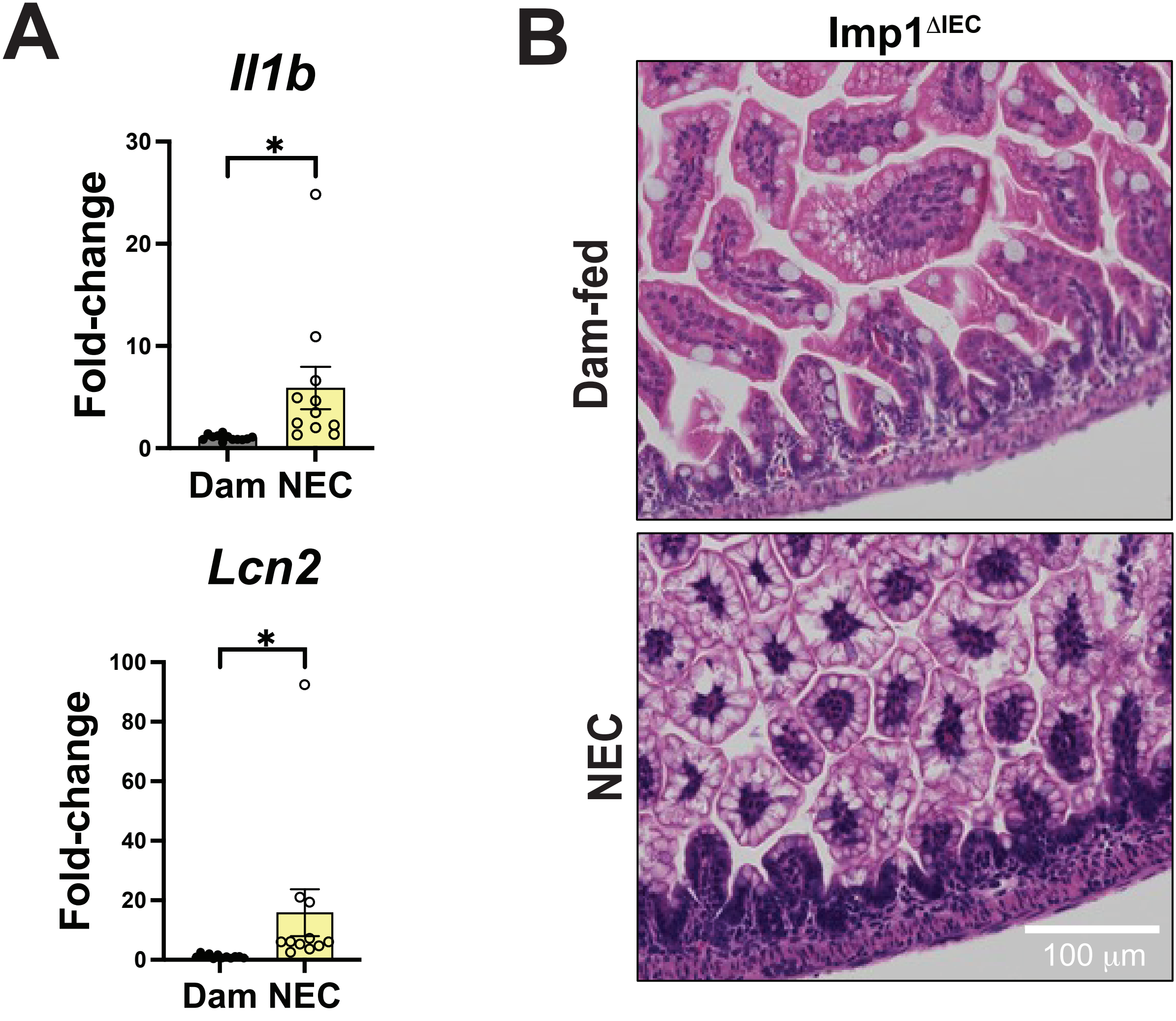


Supplemental Figure 2.

### Supplemental Figure 3

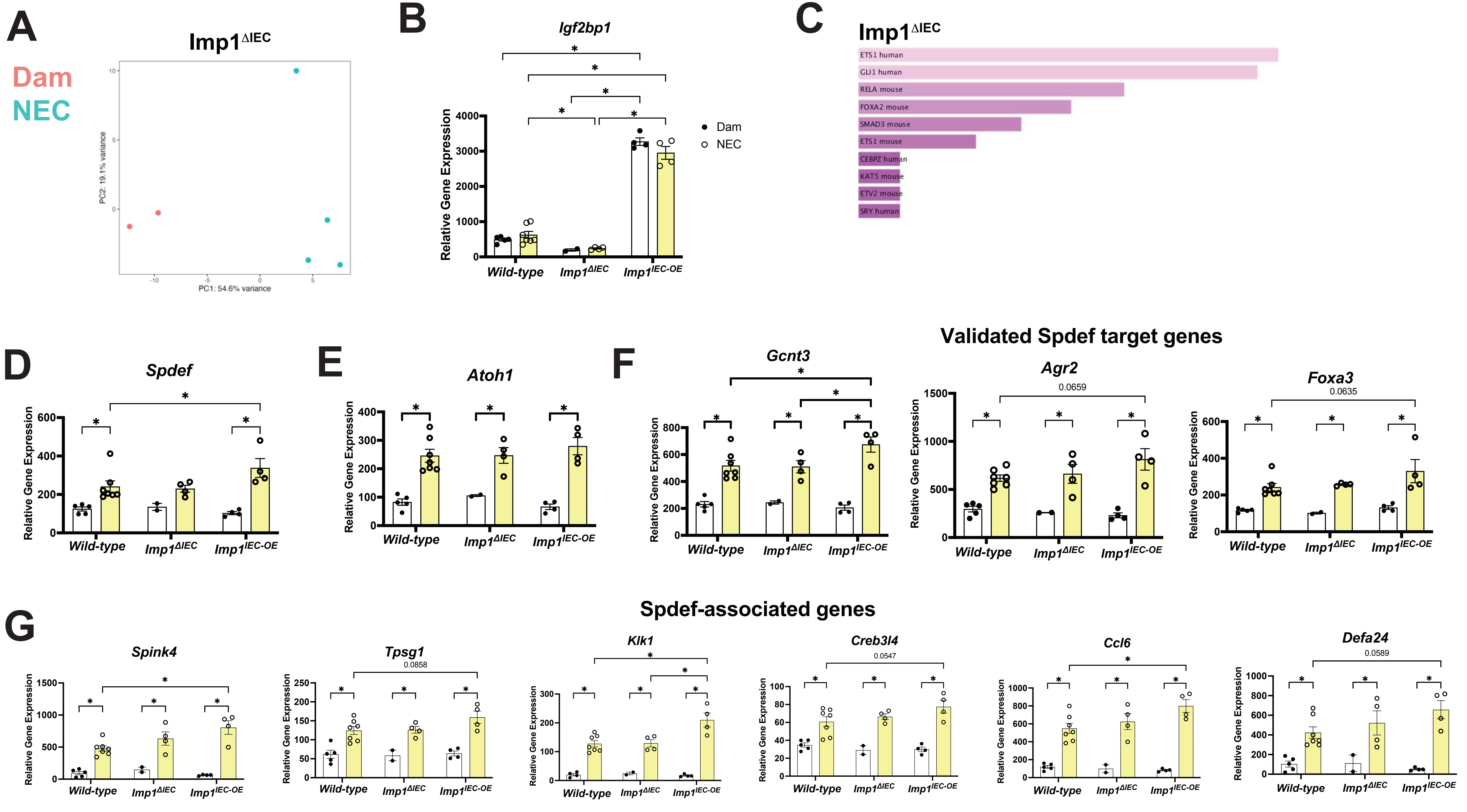


Supplemental Figure 3

### Supplemental Figure 4

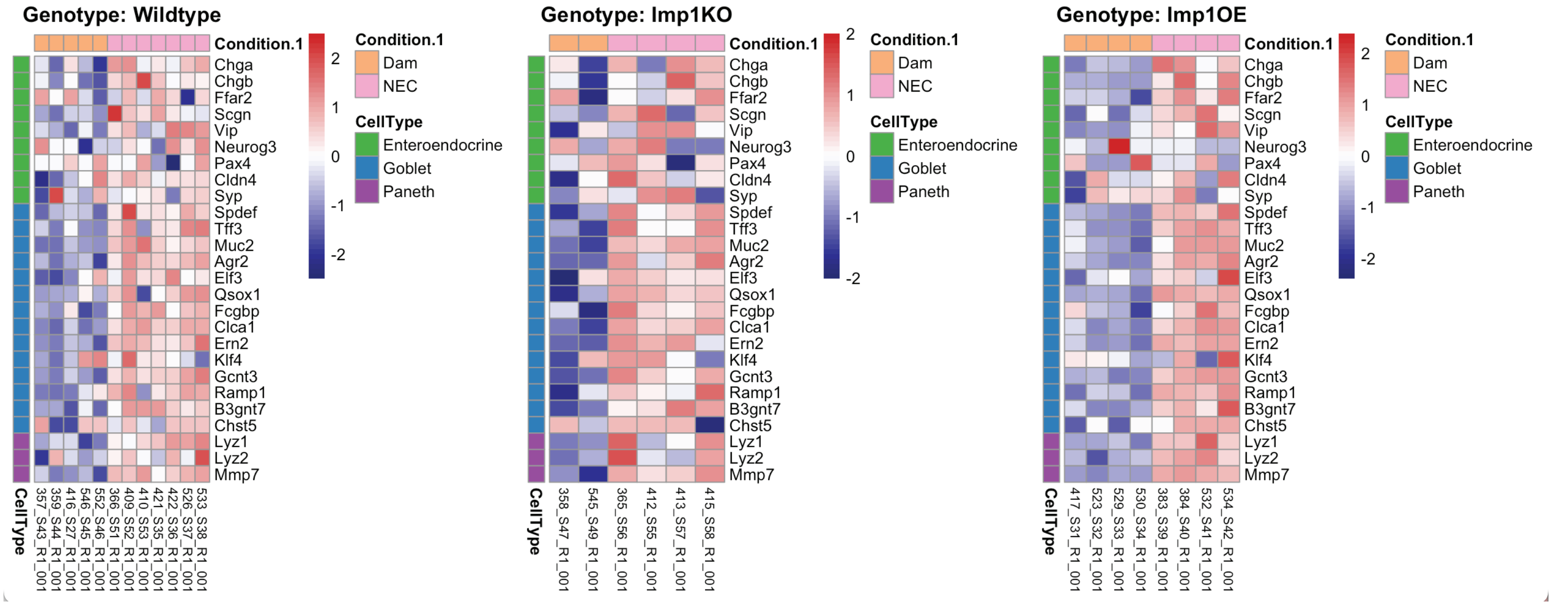


#### Supplemental Figure 4.

### Supplemental Figure 5

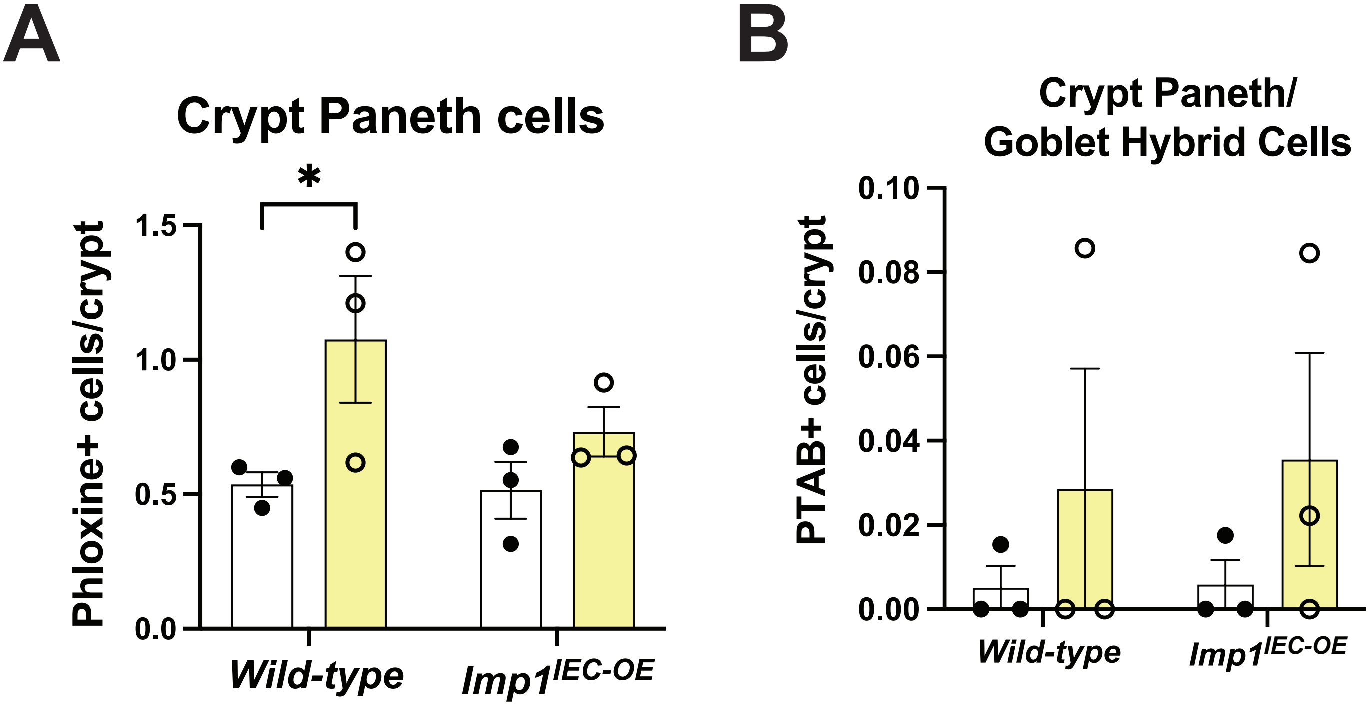


Supplemental Figure 5.

### Supplemental Figure 6

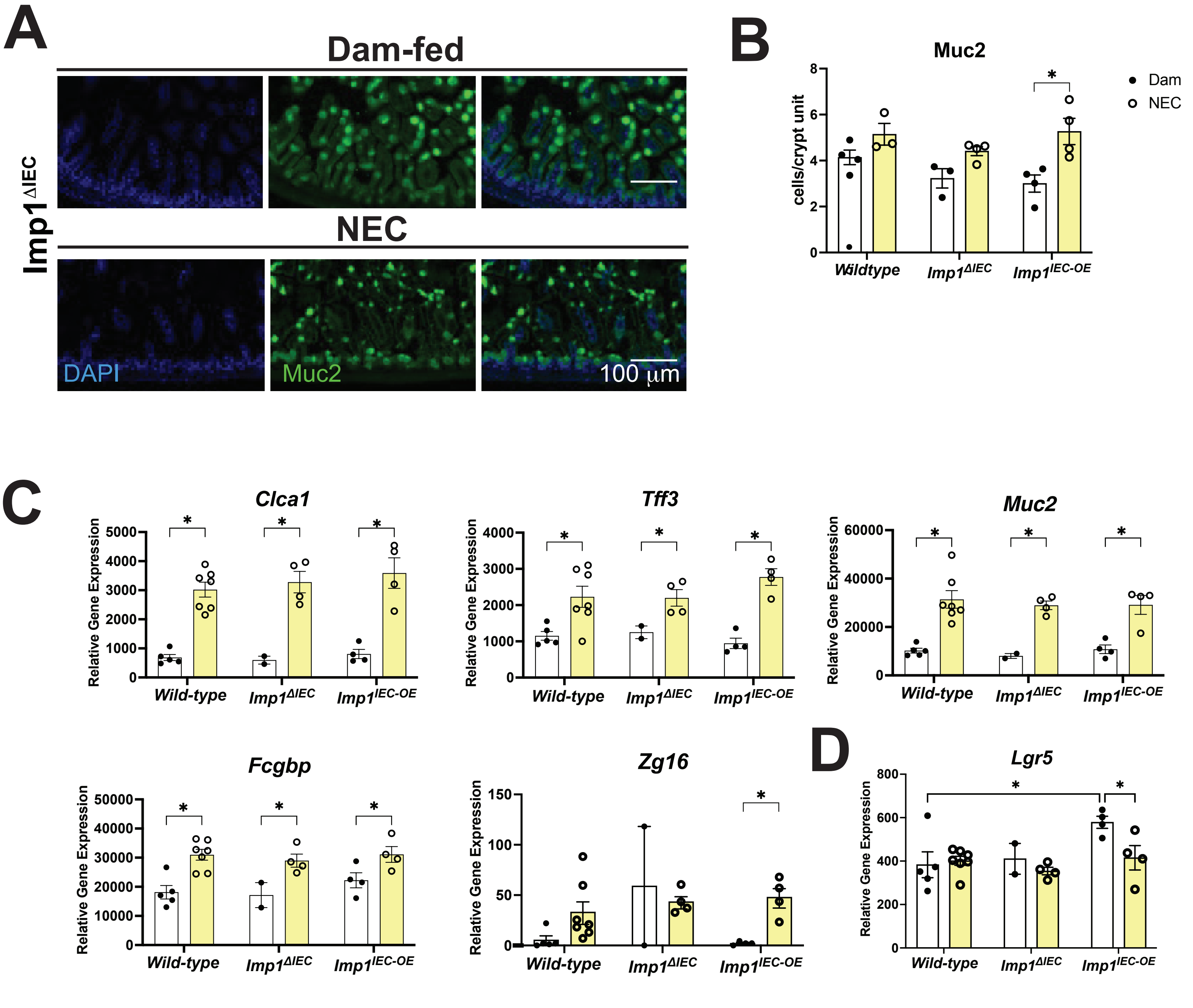


Supplemental Figure 6.

### Supplemental Figure 7

# A

##### Mucus glycosylation genes

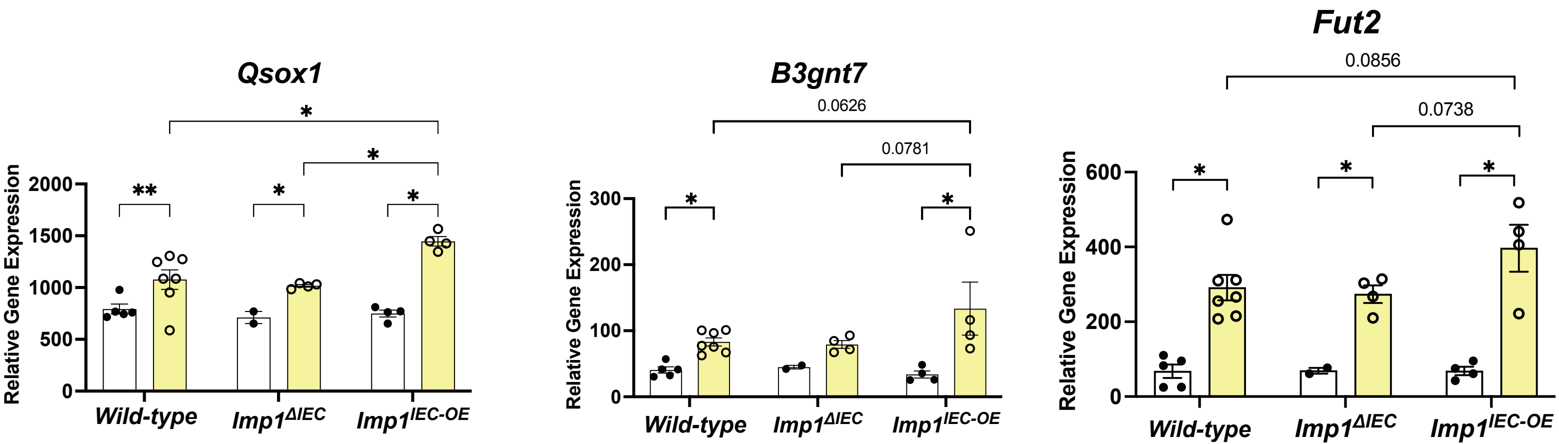


#### Supplemental Figure 7.
